# Impact of suboptimal APOBEC3G neutralization on the emergence of HIV drug resistance in humanized mice

**DOI:** 10.1101/764381

**Authors:** Matthew M. Hernandez, Audrey Fahrny, Anitha Jayaprakash, Gustavo Gers-Huber, Marsha Dillon-White, Annette Audigé, Lubbertus C.F. Mulder, Ravi Sachidanandam, Roberto F. Speck, Viviana Simon

## Abstract

HIV diversification facilitates immune escape and complicates antiretroviral therapy. In this study, we take advantage of a humanized mouse model to probe the contribution of APOBEC3 mutagenesis to viral evolution. Humanized mice were infected with isogenic HIV molecular clones (HIV-WT, HIV-45G, HIV-ΔSLQ) that differ only in their ability to counteract APOBEC3G (A3G). Infected mice remained naïve or were treated with the RT inhibitor lamivudine (3TC). Viremia, emergence of drug resistant variants and quasispecies diversification in the plasma compartment were determined throughout infection. While both HIV-WT and HIV-45G achieved robust infection, over time HIV-45G replication was significantly reduced compared to HIV-WT in the absence of 3TC treatment. In contrast, treatment response differed significantly between HIV-45G and HIV-WT infected mice. Antiretroviral treatment failed in 91% of HIV-45G infected mice while only 36% of HIV-WT infected mice displayed a similar negative outcome. Emergence of 3TC resistant variants and nucleotide diversity were determined by analyzing 155,462 single HIV reverse transcriptase (*RT*) and 6,985 *vif* sequences from 33 mice. Prior to treatment, variants with genotypic 3TC resistance (RT-M184I/V) were detected at low levels in over a third of all animals. Upon treatment, the composition of the plasma quasispecies rapidly changed leading to a majority of circulating viral variants encoding RT-184I. Interestingly, increased viral diversity prior to treatment initiation correlated with higher plasma viremia in HIV-45G but not in HIV-WT infected animals. Taken together, HIV variants with suboptimal anti-A3G activity were attenuated in the absence of selection but display a fitness advantage in the presence of antiretroviral treatment.

**IMPORTANCE:** Both viral (e.g., reverse transcriptase, *RT*) and host factors (e.g., APOBEC3G (A3G)) can contribute to HIV sequence diversity. This study shows that suboptimal anti-A3G activity shapes viral fitness and drives viral evolution in the plasma compartment of humanized mice.

## INTRODUCTION

HIV diversity is extensive on both an individual level and a global level. It drives viral adaptation in distinct cellular environments and facilitates escape from immune surveillance and antiretroviral therapy (ART,(1)). A combination of HIV features including high *in vivo* mutation rate, high replication rate and recombination between co-packaged genomes contribute to viral diversification (2, 3). Although the high mutation rate is caused by the error-prone nature of the HIV reverse transcriptase (*RT*, (4, 5)), the mutagenesis by host apolipoprotein B mRNA editing enzyme, catalytic polypeptide-like 3 (APOBEC3) cytidine deaminases also contributes to diversity (6–8). Several of the APOBEC3 family members limit HIV replication if left unchecked by HIV Vif, by mutating the viral cDNA during reverse transcription introducing guanosine to adenosine (G-to-A) substitutions in HIV provirus (9–13). Counteraction of APOBEC3G (A3G), APOBEC3F (A3F) and the stable haplotypes of APOBEC3H (A3H) by HIV Vif is essential for establishing a robust infection *in vivo* (reviewed in (10, 14)). However, proviruses in HIV infected patients frequently display high numbers of G-to-A mutations in dinucleotide contexts suggesting previous suboptimal anti-APOBEC3 activity (15–17). In fact, HIV Vif alleles obtained from the plasma and/or peripheral blood cell compartment of patients differ – to some extent – in their ability to counteract the different APOBEC3 proteins (18–20).

Our understanding of the impact of HIV Vif variation on HIV/AIDS disease outcomes remains incomplete since such studies in patients are inherently limited in scope and descriptive in nature (11, 21, 22). With the advent of the humanized mouse model in the past decade, *in vivo* studies investigating HIV pathogenesis (23–25), novel therapeutic interventions (26–29) and viral evolution (30–32) under controlled experimental conditions have become possible. Experiments in different humanized mice systems (e.g., NOG, NSG, BLT) established that HIV Vif is necessary for infection (33, 34). Moreover, the impact of A3F, A3G and A3H on replication has been tested using HIV Vif mutant viruses that are defective in counteracting A3D, A3F, A3G or A3H (35–37). These studies show that the failure to neutralize A3G results in severe attenuation of viral replication with viruses unable to counteract A3G being less diverse than their wild-type counterparts (35, 36). Thus, interrogation of A3G-driven HIV diversification in an “all or nothing” fashion indicates that a complete loss of anti-A3G activity results in HIV restriction and limits viral diversification.

However, most circulating HIV strains maintain some activity against A3G such that experiments with viruses with suboptimal anti-A3G activity may provide a more relevant picture regarding the effects of partial APOBEC3 neutralization. Moreover, we reasoned that to directly test to what extent variation in APOBEC3 neutralization capacity influences HIV evolution *in vivo*, one needs to perturbate not only the HIV Vif-A3G axis (e.g., by using HIV Vif mutants) but also the viral equilibrium reached upon establishment of infection (e.g., by administrating an antiretroviral to apply selection pressure). Thus, in this study, we infected humanized mice with wild-type and selected Vif mutant viruses and monitored infection in the plasma compartment over time in the presence and absence of antiretroviral treatment in the form of 3TC monotherapy. We assessed viral replication by longitudinally quantifying plasma viremia as well as viral diversification using a high-resolution, molecular ID tag based deep sequencing approach. Our data indicate that suboptimal neutralization of A3G results in attenuated viral replication in the absence of selection but provides a replication advantage in the presence of 3TC antiretroviral treatment.

## MATERIALS AND METHODS

### Ethics statement

Animal experiments were approved by the Cantonal Veterinary Office (#26/2011 & #93/2014) and performed in accordance to local guidelines and to the Swiss animal protection law. The Ethical Committee of the University of Zurich approved the procurement of human cord blood and written informed consent was provided prior to the collection of cord blood.

### Cell-lines

HEK293T cells were obtained from ATCC (CRL-3216) and TZM-bl reporter cells were obtained from the AIDS Research and Reference Reagent Program, Division of AIDS, NIAID, National Institutes of Health (NIH AIDS Reagent Program, cat. 8129) (38–42). HEK 293T cells and TZM-bl reporter cells were maintained in Dulbecco’s modified Eagle medium (DMEM, Fisher Scientific, cat. MT10-013-CV) supplemented with 10% fetal bovine serum (FBS) (Gemini Bio-Products) and 100 U/mL penicillin-streptomycin (Fisher Scientific, cat. MT30002CI). HEK 293T and TZM-bl cells were grown on 100mm Falcon^TM^ Standard Tissue Culture Dishes (Fisher Scientific, cat. 08-772E).

### Generation of viral stocks

Isogenic molecular clones were derived from pNL4-3 (HIV-WT; NIH AIDS Reagent Program, cat. 114) (43), Replication competent molecular clones encoding Vif mutants E45G (HIV-45G) and SLQ144AAA (HIV-ΔSLQ) were generated as previously described (44). Viral stocks were generated by transfecting HEK293T cells using 4μg/mL polyethylenimine (PEI, Polysciences Inc., cat. 23966). Culture supernatants were collected 48 hours post-transfection, filtered and frozen at −80°C until further use. Viral stock concentrations were quantitated using an in-house p24 ELISA (45) and/or tittered on TZM-bl reporter cells as previously described (44).

### Generation of humanized mice

Animals were housed under specific pathogen free conditions. Humanized mice were generated as previously described (29). Briefly, newborn immunodeficient NOD-scid IL-2Rγ-null (NSG) mice (Jackson laboratory, Bar Harbor, ME) were irradiated 1-3 days after birth with 1 Gy and transplanted intrahepatically with approximately 2.0±0.5×10^5^ cord blood-derived CD34+ cells. Between 2 and 6 mice were transplanted with cells from the same donor. A total of 12 donors were used for the three infection experiments.

Twelve to sixteen weeks after transplantation, human engraftment and *de novo* human immune system reconstitution in the mice were assessed by staining peripheral blood with monoclonal antibodies against the panhuman marker CD45 (Beckman Coulter, cat. B36294), CD19 (Biolegend, cat. 302212), CD3 (Biolegend, cat. 300308), CD4 (Biolegend, cat. 300518) and CD8 (Biolegend, cat. 301035). Flow-cytometry analyses were performed on a CyAN™ ADP Analyzer (Beckman Coulter, Brea, CA).

### Infection and antiretroviral treatment of humanized mice

Mice were infected intraperitoneally with 2×10^5^ TCID50 per mouse of each of the 3 HIV clones in 200μL volume. Plasma viremia was measured using Cobas® Amplicor technology (Roche, Switzerland) at the described time points throughout the infection. The detection limit of the assay is 400 HIV RNA copies/mL.

Lamivudine (3TC Epivir, GlaxoSmithKlein, UK) treatment was started 30 days post infection in the treatment group of the infected mice. 3TC tablets were weighed, pulverized and combined with food pellets as previously described (29).

### Amplification of HIV from plasma viral RNA

Viral RNA was extracted from 140μL frozen plasma using the QIAamp Viral RNA Minikit (QIAGEN, cat. 52904) as per manufacturer’s instruction.

For deep sequencing, cDNAs were synthesized using custom reverse transcription primers.(Supplemental Table 1; Integrated DNA Technologies). From 5’ to 3’, our first generation *RT* cDNA primer (4372) included a 16 basepair (bp) HIV-specific sequence (accession: AF324493.2: 3336-3351), 8 bp randomized sequence (unique molecular ID (UMID)) and an additional 23 bp HIV-specific sequence (AF324493.2: 3305-3327). Viral cDNA was synthesized using the Invitrogen^TM^ ThermoScript RT-PCR System (Thermo Fisher Scientific, cat. 11146016). Briefly, RNA and primer were denatured at 65°C for 5 min. ThermoScript reaction mix was added to the RNA and primer and incubated at 50°C for 60 min, followed by an inactivation step of 85°C for 5 min.

Viral cDNA was column purified (Zymo Clean and Concentrator kit, cat. D4034) and amplified in a first round PCR using primers 1922 (AF324493.2: 2929-2946) and 1923 (AF324493.2: 3337-3356). First round PCR used Pfx50 polymerase (94°C for 2 min, followed by 23 cycles of 93°C for 15 sec, 48°C for 30 sec, 68°C for 60 sec and a final extension of 68°C for 10 min) (Thermo Fisher Scientific, cat. 12355012). First round products were purified (Zymo Clean and Concentrator) and a second PCR was performed to add Illumina-based adapters using custom primers 1690 and one of 73 primers with distinct MiSeq barcode identifiers (Supplemental Table 1). The second round PCR used PfuUltra II Fusion HS Polymerase (95°C for 2 min, followed by 25 cycles of 93°C for 20 sec, 50°C for 20 sec, 72°C for 15 sec and a final extension of 72°C for 10 min) (Agilent Technologies, cat. 600672). Second round PCR products were confirmed by electrophoresis and purified by SPRIselect magnetic bead selection (Beckman Coulter, cat. B23317).

To amplify *vif* sequences from plasma vRNA, a *vif* specific cDNA synthesis primer was used (4373). From 5’ to 3’, primer 4373 included a 16 bp T7 sequence, an 8 bp randomized sequence and 23 bp HIV-specific sequence (AF324493.2: 5386-5406). Viral cDNA was synthesized using the ThermoScript RT-PCR System and purified as described above. cDNAs were amplified in the first round PCR using primers 2402 (AF324493.2: 4993-5012) and 2401 (T7). First round products were purified and amplified in a second round PCR using primers 2403 and one of 80 primers with distinct MiSeq barcode identifiers and complementarity to the T7 sequence. PCR cycling conditions were the same as used for the *RT* products.

### MiSeq Library Preparation and MiSeq Instrumentation

Sequencing libraries were run on the Illumina MiSeq to sequence paired-end reads. To prepare libraries, bead-purified PCR products containing Illumina adapters (376 bp *RT* amplicons and 393 bp *vif* amplicons) were quantitated by Qubit dsDNA HS Assay Kit (Thermo Fisher Scientific, cat. Q32854). To sequence 250 bp of the RT region, a 6pM final library was run with a 20% spike-in of PhiX Control V3 (Illumina, cat. FC-110-3001) for 2×150 cycles using MiSeq Reagent Kit v2 (300 cycles, Illumina, cat. MS-102-2002) or MiSeq Reagent Kit v3 (600 cycles, Illumina, cat. MS-102-3003). Custom sequencing primers were used for the forward (1692), index (3890), and reverse (3889) reads. To sequence a 269 basepair long region of *vif*, a 6pM final library was run with a 15% spike-in of PhiX Control V3 for 2×150 cycles. Custom sequencing primers were used for the forward (4580), index (4577) and reverse (4578) reads (Supplemental Table 1).

### Bioinformatic Pipeline for Sequence Analyses

Custom Unix and Perl scripts were written to process FASTQ files. First, paired-end reads were merged using Paired-End reAd mergeR (46). Reads were aligned to pNL4-3 *RT* or *vif* reference sequences (AF324493.2) and filtered. Sequences were grouped by distinct UMIDs. A minimum of three high quality merged pair-end reads contained the same UMID were required to generate a consensus sequence. Consensus sequences were defined as sequences where each nucleotide reflected 70% or more of all nucleotides sequenced for that given position. Consensus sequences for sequences defined by a distinct UMID were identified using in-house scripts. Twenty-five UMIDs that differed by only one mismatch from the HIV template sequence were considered to be experimental artifacts and excluded from the analyses.

Additional custom scripts were written to identify 3TC resistance mutations at *RT* codon 184 and to compute overall mutation rates, GG-to-GA as well as GA-to-AA mutagenesis and the frequency of stop codons within the sequenced *RT* and *vif* regions. DNASP v5 polymorphism software was used to calculate the nucleotide diversity (π), defined as the average pair-wise number of nucleotide differences per site in all possible pairs of consensus sequences per sample (47).

### Statistics

Normality was assessed using the D’Agostino and Pearson test for numerical data such as baseline plasma viremia, changes in viremia, proportions of 3TC susceptible/resistant viral sequences, nucleotide diversity, and mutation rates. If groups passed normality tests, parametric student’s t-test for unpaired data or paired student’s t-test for paired data were used. Otherwise, non-parametric Mann-Whitney or Wilcoxon matched pairs signed-rank tests were used. For categorical data (e.g., treatment failure/success), Fisher’s exact test was used. Linear regression models were used for replication kinetics and F-tests were used to compare slopes of curves of best fit. Finally, nonlinear regression methods (exponential (Malthusian)) using weighted least squares were used with extra sum-of-squares F-test to compare nonlinear curves of best fit.

Significance testing is reported as exact p-values in the text or using asterisks (*, p ≤ 0.05; **, p ≤ 0.01; ***, p ≤ 0.001; ****, p ≤ 0.0001). All statistics were performed using the built-in analysis packages from GraphPad Prism v7.0 Suite (GraphPad Software, Inc, La Jolla, CA).

### Data availability

All processed FASTA sequencing files are publicly available (will be released upon publication). In addition, in-house codes for sequence read processing can be accessed at https://github.com/AceM1188/Vif-pipeline.

## RESULTS

### HIV-WT and HIV-45G establish productive infection in the humanized mouse model system

We performed three independent long-term infection experiments in 48 humanized NSG (hu-NSG) mice mimicking natural infection (Exp. 1; 85 days follow-up post infection) or treatment interventions (Exp. 2 and Exp. 3; 58-75 days of follow up post infection). Fig. 1A depicts the time line for each experiment including the viruses used for infection and the time points at which blood samples were collected for further analysis (e.g., plasma viremia, HIV RNA sequence analysis).

**Fig. 1:**
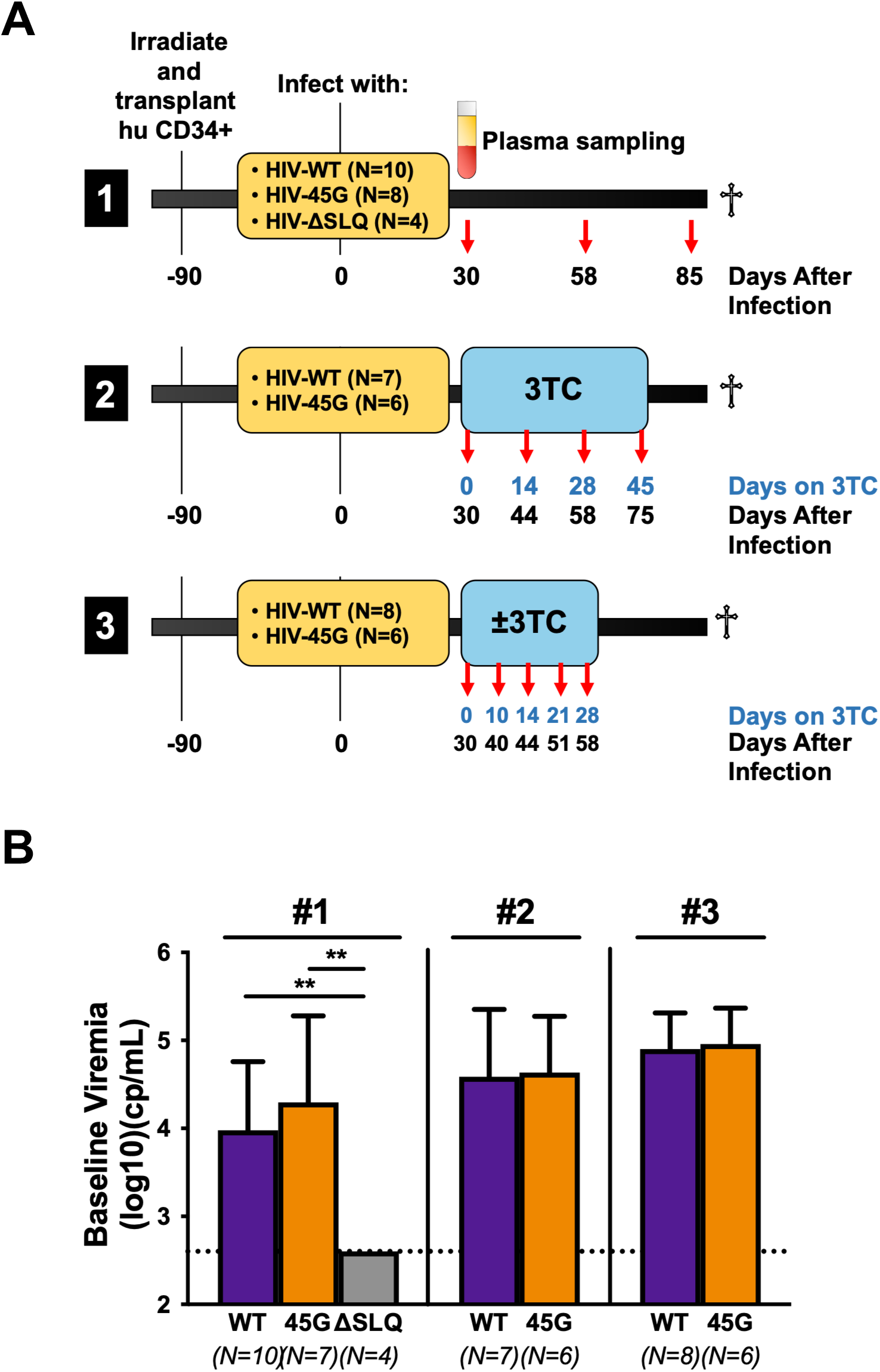
Infection of humanized mice with HIV-Vif variants. **(A)** Newborn NOD-scid IL-2Rγ-null (NSG) mice were irradiated after birth and transplanted with human donor cord blood-derived CD34+ cells. Mice were infected intraperitoneally with 2×10^5^ TCID_50_ of virus (HIV-WT, HIV-45G or HIV-ΔSLQ). The specifics for each of the three infection experiments are provided in the time-lines. Plasma was collected at the indicated time points for viremia measurements and sequence analysis. **(B)** Comparison of the plasma viremia Day 30 post infection (baseline) in mice infected with the different viruses in the three different experiments. The lower limit of detection of the assay is 400 copies/mL (cp/mL). The mean and standard deviations of the viral loads are depicted. Viremia for HIV-ΔSLQ is significantly different from HIV-WT or HIV-45G (p ≤ 0.0020, Mann-Whitney).

The three viruses selected for these experiments (HIV-WT, HIV-45G and HIV-ΔSLQ) consist of isogenic clones of the HIV NL4-3 isolate that differ only in their ability to counteract A3G. These mutants have been well characterized previously by us and others (12, 18, 44). Briefly, NL4-3 serves as HIV-WT. NL4-3 Vif counteracts A3G, A3F and A3D but is inactive against the stable A3H haplotypes (48). HIV-ΔSLQ fails to bind Cullin5 E3 ligase complex due to three alanines in place of the SLQ BC-box motif (144–146) and, therefore, fails to counteract any of the APOBEC3 proteins. HIV-45G carries a single point mutation in codon 45 of Vif (HIV-45G) (18, 44, 49). This mutation in Vif results in attenuation but not complete abrogation of its activity against A3G while preserving activity against A3F and A3D (18). Importantly, residue 45 of Vif has not been associated with other Vif functions and is not involved in any other viral gene products (50, 51). HIV-WT and HIV-45G replicate to comparable levels in primary human peripheral mononuclear cells while HIV-ΔSLQ fails to initiate a spreading infection in cell culture (44). Importantly, suboptimal neutralization of A3G by HIV-45G does not result in a sizeable replication defect in short-term cell culture infection experiments making it very well suited for *in vivo* infection experiments aimed at studying evolution in humanized mice.

Hu-NSG mice were reconstituted with CD34+ cells isolated from umbilical cord blood and immune reconstitution was confirmed after 90 days prior to infection. In all three experiments, we measured viremia at Day 30 post infection. Infection with HIV-WT and HIV-45G established comparable average viremia with no significant difference between the three independent infection experiments (Fig. 1B). In good agreement with previous findings, a functional HIV Vif was necessary to establish a productive infection since all four hu-NSG mice infected with HIV-ΔSLQ displayed no detectable plasma viremia at Day 30 post infection (Fig. 1B).

### Replication of HIV-45G but not HIV-WT is attenuated over time in humanized mice

In the natural infection experiment (Exp. 1), we followed viral replication in the plasma compartment for nearly three months post infection. Plasma viremia was measured using molecular diagnostics at regular intervals. All productively infected animals (HIV-WT, N=14; HIV-45G, N=8) maintained viremia above the limit of detection until the end of the experiment. HIV-ΔSLQ infected animals never displayed viral load measurements above 400 copies/ml (limit of detection of the assay). The change in plasma viremia over time is plotted in Fig. 2A. While HIV-WT infection resulted in an increase of replication over time in all but two animals, HIV-45G replication was attenuated in the majority of infected animals within 40-60 days post infection. The change in plasma viremia between baseline (i.e., Day 30 post infection) and endpoint was significantly different between the two viruses (HIV-45G: 0.77 log_10_ *decrease* versus HIV-WT 0.43 log_10_ *increase*; p=0.0026). Taken together, *in-vivo* HIV-45G replication appears to be attenuated, although this trait becomes apparent only 50-60 days post infection.

**Fig 2:**
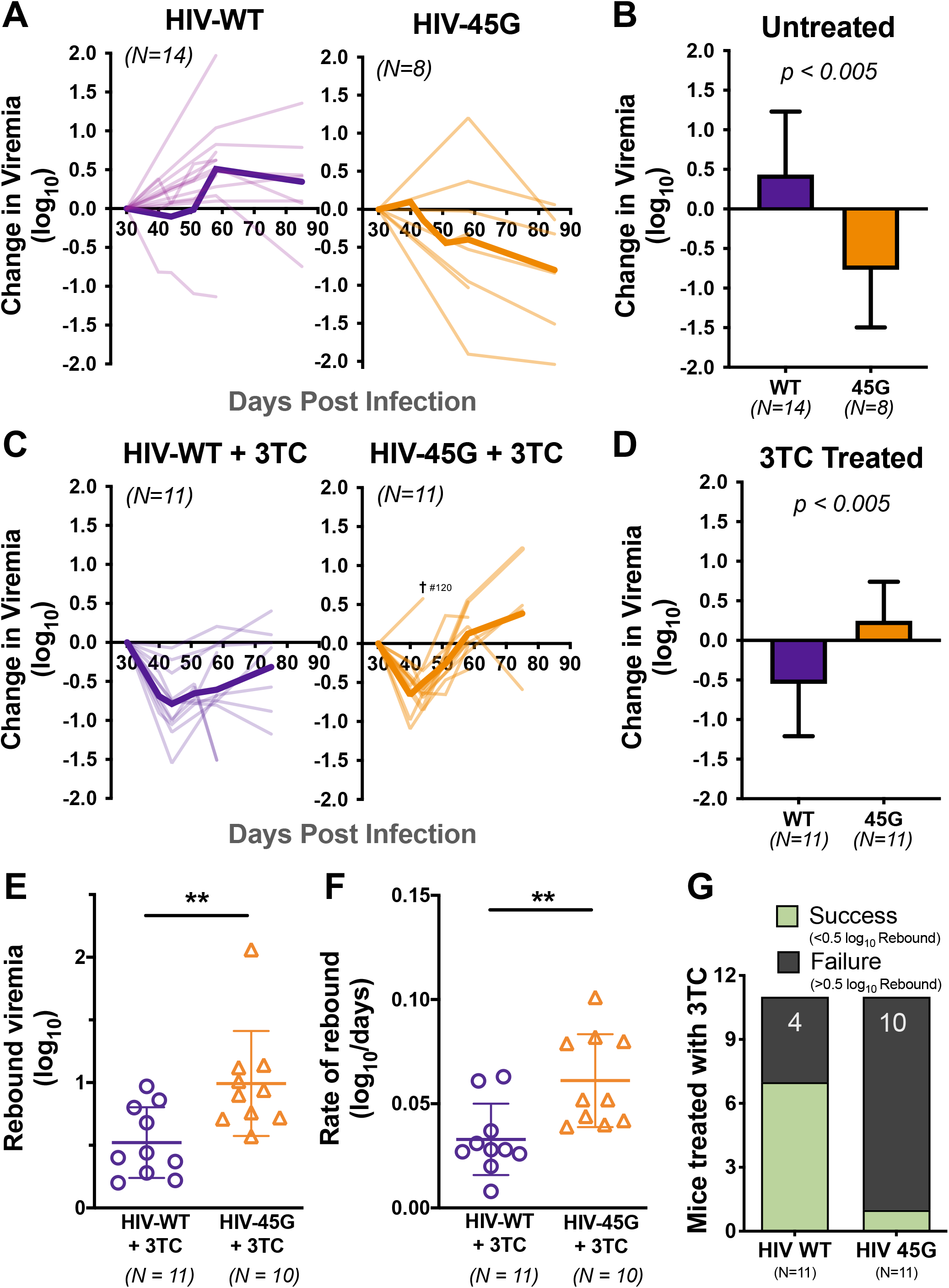
Viral replication in humanized mice in the absence or presence of 3TC treatment. **(A)** Spaghetti plots depicting the change in viremia relative to baseline viral load (Day 30 post infection) of individual untreated mice (pale lines) and mean change in viremia in untreated mice (bold lines). **(B)** Barplots depicting the change in viremia at the end of infection (last available timepoint) versus baseline viremia in untreated mice. Means and standard deviation are depicted. p = 0.0026 by unpaired student’s t-test. **(C)** Spaghetti plots depicting the change in viremia in 3TC treated mice. **(D)** Barplots depict the overall change in viremia at the end of infection in 3TC treated mice with means and standard deviations. p = 0.0045 by unpaired student’s t-test. **(E)** Rebound viremia – defined as the maximum fold rebound viral load from nadir (maximum level of suppression observed; log(VL_max_/VL_nadir_) – is depicted for mice treated with 3TC. Each point represents an individual mouse. Mean and standard deviations are depicted. p = 0.0068 by Mann-Whitney test. **(F)** Rate of rebound was also measured over the time from nadir to maximum viremia. Means and standard deviations depicted. p = 0.0052 by unpaired student’s t-test. **(G)** Qualitative assessment of treatment outcomes. 3TC treatment was defined as successful if the viral rebound was less than 0.5 log_10_ from nadir. Conversely, treatment failure was met if the viral rebound was greater than 0.5 log_10_ from nadir. p = 0.0237, Fisher’s exact test.

### Suboptimal neutralization of A3G confers superior viral fitness in the presence of 3TC in the humanized mouse model system

In the treatment intervention infection experiments (Exp. 2 and Exp. 3), animals robustly infected with either HIV-WT (N=11) or HIV-45G (N=11) were treated with 3TC starting at Day 30 post infection until the end of infection (e.g., Day 75 in Exp. 2 and Day 58 in Exp. 3). Of note, the blood collection intervals in Exp. 3 were shorter than in Experiment 2 (4-7 day versus 14 day intervals).

The initial virological response upon 3TC initiation was comparable between the two groups. Within the first two weeks of 3TC treatment both HIV-WT and HIV-45G viremia decreased to comparable nadirs (0.69 log_10_ versus 0.66 log_10_, respectively) (Fig. 2C). One mouse (#120, infected with HIV-45G) was euthanized at Day 44 post infection as per animal safety protocol due to signs of wasting. Considering the overall change from baseline to experiment termination, HIV-WT replication *decreased* by 0.55 log_10_ in the presence of 3TC, whereas HIV-45G viremia had *increased* by 0.25 log_10_ from baseline (p=0.0045) (Fig. 2D). Moreover, HIV-45G infected animals experienced a larger (0.99 log_10_ vs. 0.52 log_10_, p=0.0068, Fig. 2E) and faster (0.061 vs. 0.033 log_10_/day, p=0.0052, Fig. 2F) viral rebound from the initial lowest points reached upon 3TC treatment initiation.

We also assessed 3TC treatment outcomes in a qualitative manner (e.g., treatment success versus failure, with success being defined as sustained reduction in viremia with less than 0.5 log_10_ rebound from the lowest points). Antiviral treatment was successful in 64% of HIV-WT infected mice but only 9% of HIV-45G infected mice (Fig. 2G). Thus, HIV-45G infected mice had a significantly higher risk of failing treatment (RR: 7.000; 95% CI 1.482 – 40.54; p=0.0237).

Taken together, the replication of HIV-45G in the absence (Exp. 1) and in the presence of 3TC (Exp. 2 and Exp. 3) was very different (compare Figs. 2B and 2D). While the fitness of HIV-45G was attenuated over time in naïve animals, its replication was significantly less affected by 3TC than that of HIV WT suggesting the existence of a selection advantage.

### Dynamics of genotypic 3TC drug resistance

The molecular mechanisms resulting in 3TC resistance are well described (52–58). Single point mutations in codon 184 of HIV *RT* emerge rapidly upon 3TC treatment both *in vivo* and in cell culture. RT-184I (AT<u>G</u>->AT<u>A</u>) and RT-184V (<u>A</u>TG-><u>G</u>TG) are the most common substitutions observed in HIV infected patients failing 3TC-containing ART (59) (Fig. 3A). These substitutions confer up to >1,000-fold reduced susceptibility to 3TC (53, 58). Both mutations can result from reverse transcription errors, although the mutation leading to RT-184I is also within a dinucleotide context favored for A3G driven mutagenesis (e.g., GG-to-AG mutations, (AT<u>G</u>G->AT<u>A</u>G))(55, 60).

**Fig. 3:**
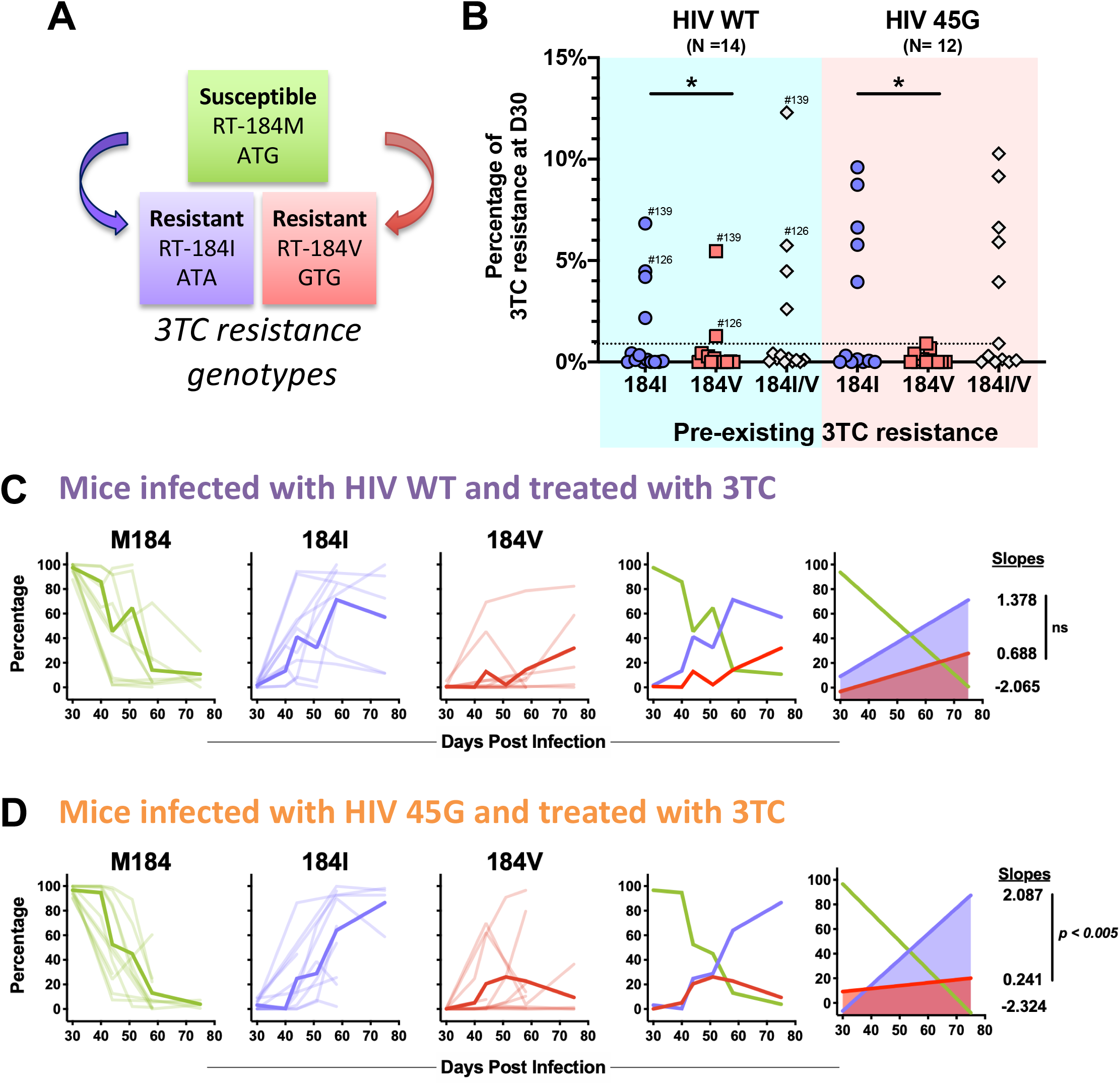
Drug resistance development in 3TC treated mice. **(A)** Genotypic 3TC drug resistance is due to single point mutations in codon 184 of HIV reverse transcriptase (*RT)* gene. Methionine (M184) represents the susceptible wild-type sequence while RT-184I or RT-184V render the virus resistant to 3TC. **(B)** Pre-existing 3TC resistance detected at Day 30 post infection (”D30”) prior to 3TC treatment initiation. Each dot represents the percentage of viruses encoding RT-184I or RT-184V in a given mouse. Minority 3TC resistant viral populations were defined as representing at least 1% of the total number of UMIDs sequenced for each mouse at this time point (dotted line). Some mice harbored both M184I/V variants and are indicated by mouse ID number. (*) p ≤ 0.05, Wilcoxon matched pairs signed rank test. **(C)** Spaghetti plots longitudinally depicting the relative proportion of 3TC susceptible and resistant viral variants in HIV-WT infected mice. M184, RT-184I and RT-184V data for individual mice (pale lines) and mean proportions at each timepoint (bold lines) are depicted. Slopes of lines of best fit were calculated to measure kinetics of RT-184M, RT-184I and RT-184V variants. Slopes compared by F-test (ns: not significant, p = 0.5372). **(D)** Spaghetti plots longitudinally depicting the relative proportion of 3TC susceptible and resistant viral variants in HIV-45G infected mice. Slopes compared by F-test (p = 0.0035).

To further investigate the *in-vivo* dynamics of resistance appearance, we used a next generation sequencing approach to analyze a 250 bp region of HIV *RT* (codon 177 to 258) from cell-free HIV RNA (vRNA) present in the plasma compartment. We combined a unique molecular ID (UMID) strategy, to compensate for PCR-mediated amplification bias and errors (61, 62), with 150 bp paired-end Illumina sequencing chemistry. Briefly, two series of custom primers with an 8 bp randomized IDs were used to tag HIV vRNA during the reverse transcription step prior to amplification and sequencing. We sequenced a total of 111 vRNA samples obtained from 34 animals (primer #4372: 69 vRNA, 14 mice; primer #4633: 42 vRNA, 20 mice) generating a total of 9,705,468 high-quality paired-end reads representing 155,462 UMIDs (=individual HIV genomes). On average, we obtained 1,400 UMIDs for each individual plasma sample generating between 1,333 and 17,854 unique *RT* consensus sequences for each infected animal over the course of the infection.

This approach, importantly, provides sufficient resolution to identify minority viral populations and provide insights into the composition of the viral quasispecies. We first analyzed the mutations present at codon 184 of RT (Figs. 3B-3D). Minority 3TC resistant populations (defined as 1% or more of the overall number of UMIDs present in a given sample) were detectable after Day 30 post infection but prior to 3TC treatment in a third of the HIV-WT (N=4) and HIV-45G (N=5) infected mice (“pre-existing 3TC drug resistance” Fig. 3B). RT-184I resistant viruses were far more common than RT-184V (HIV-WT, p=0.0342; HIV-45G, p=0.0273) but generally made up less than 10% of the sampled circulating viruses in a given animal. A combination of RT-184V and RT-184I was found in two animals (identified by the specific mouse number in Fig. 3B). Of note, when we stratified by treatment outcome, pre-existing 3TC drug resistance was not associated with treatment failure (p=0.5227, Fisher’s exact test) and did not display any significant relationship with rate of nadir formation (HIV-WT, p=0.0762; HIV-45G, p=0.1115) or rebound rate (HIV-WT, p=0.3267; HIV-45G, p=0.0849).

We next examined the dynamics of 3TC drug resistance in treated mice (Figs. 3C-3D). In most animals, the 3TC-susceptible RT-184M majority was rapidly replaced with viruses encoding 3TC-resistant RT-184I or RT-184V alleles. On rare occasions, substitutions other than Valine or Isoleucine were detected at codon 184 (e.g., ACG (T), AAG (K)). The kinetics of RT-184I and RT-184V appearance in the plasma compartment were comparable in HIV-WT infected mice (p=0.2107, Fig. 3C) but RT-184I variants emerged 8.7 times more rapidly than RT-184V in HIV-45G infected mice (p=0.0035, Fig. 3D). Overall, the relative proportion of the HIV-45G variants with RT-184V remained stable or declined over time while HIV-45G variants with RT-184I steadily increased (Fig. 3D). Thus, emergence of viral variants with RT-184I is favored over that of RT-184V variants in animals infected with a virus displaying suboptimal A3G neutralization activity. Of note, M184I variants also appear often prior to M184V variants in 3TC treated patients (53, 58).

### Characterization of *RT* sequence diversity throughout the course of infection

We next determined sequence diversity within the sequenced RT region beyond the 3TC drug resistance associated codon 184. We calculated the nucleotide diversity (π) among unique RT variants within a given plasma sample (63–65).

Prior to 3TC treatment (Day 30 post infection), mice infected with HIV-WT and HIV-45G showed comparable nucleotide diversity in RT (p=0.6308, Fig. 4A). However, when we assessed the relationship between RT nucleotide diversity and the plasma viral load measured at Day 30 (prior to treatment initiation), we noted that nucleotide diversity positively correlated with the level of plasma viremia in HIV-45G infected animals (Fig. 4C) while the opposite was true for HIV-WT infected animals (Fig. 4B). For this analysis we assumed an exponential (Malthusian) relationship based on the nature of HIV growth and diversity in acute infection (66, 67).

**Fig. 4:**
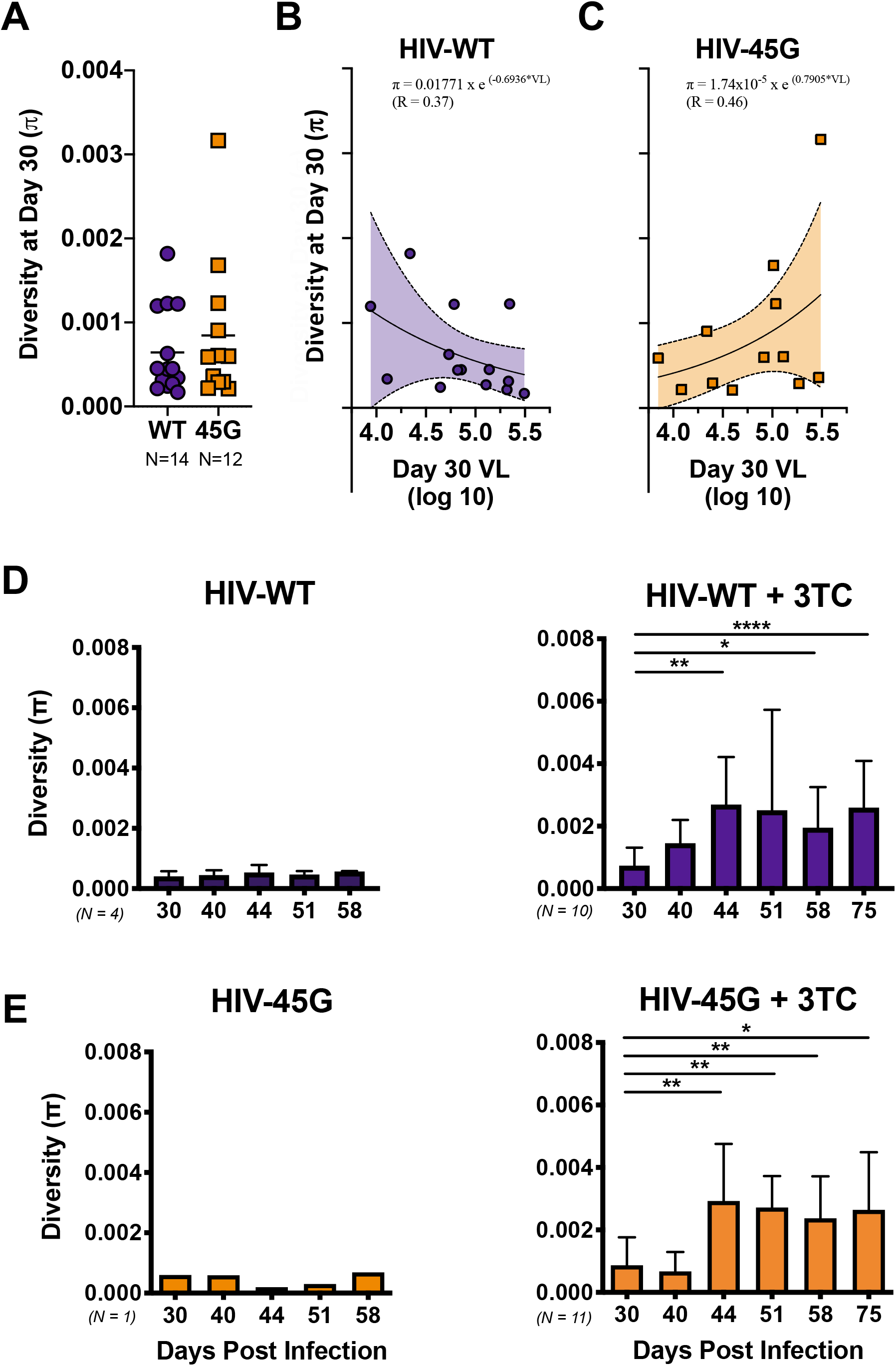
Genetic diversity of circulating viruses in the plasma of infected mice. **(A)** Dot plot depicting nucleotide diversity (π) in the sequenced HIV *RT* gene in HIV-WT and HIV-45G infected mice prior to initiation of 3TC treatment (30 Days post infection). Points represent π in each mouse and bars depict means. **(B)** Diversity and viremia (VL) data in mice infected with HIV-WT prior to treatment were fit to a weighted nonlinear exponential growth (Malthusian) model. Best fit curve and 95% confidence interval (CI) bands are portrayed. The corresponding curve equation and weighted correlation coefficient (R) are depicted above. **(C)** Diversity and VL data in mice infected with HIV-45G prior to treatment fit to an exponential growth model as in (B). **(D)** Diversity in plasma viruses of HIV-WT infected mice over time (untreated left, 3TC treated right). Significance determined by Mann-Whitney test (*, p ≤ 0.05; **, p ≤ 0.01; ***, p ≤ 0.001; ****, p ≤ 0.0001). **(E)** Diversity in plasma viruses of HIV-45G infected mice over time (untreated left, 3TC treated right). Significance determined by Mann-Whitney test (*, p ≤ 0.05; **, p ≤ 0.01; ***, p ≤ 0.001; ****, p ≤ 0.0001).

We next explored how RT sequence diversity changed throughout the course of infection. While RT diversity remained largely unchanged in treatment-naïve animals, both HIV-WT and HIV-45G diversity in treated animals initially increased but then stabilized (Day 44 post infection, Fig. 4D-E). RT sequence diversity in 3TC treated mice was driven by drug resistance associated mutations. Indeed, mutations at *RT* codon 184 contributed to 49-74% of HIV-WT and 45-69% of HIV-45G diversity (data not shown). When we excluded codon 184 from π analyses, any increases in diversity from baseline to Day 44 through Day 75 were lost in the HIV-WT infected mice (p≥0.3097). However, when we did the same for HIV-45G infected mice, HIV-45G viruses displayed a significant increase in diversity starting at Day 58 through Day 75 post infection (p≤0.0430, Mann-Whitney). Thus, while the observed increase in *RT* sequence diversity is mainly due to selection of 3TC resistant variants, other sites within RT contribute to viral diversity in HIV-45G infected mice. However, these non-drug resistance associated changes in HIV-45G viruses are only observed at later time points suggesting that they require more time to appear.

Next, we explored the contribution of APOBEC3-driven mutagenesis to the *RT* sequence diversity. Towards this end, we measured G-to-A mutations within APOBEC3-specific dinucleotide motifs (e.g., A3G: GG-to-AG; A3D/A3F: GA-to-AA, (68)). The 250 bp long region of RT that we sequenced contains 15 GG and 25 GA dinucleotides that could serve as APOBEC3 target motifs. At Day 30 post infection (prior to 3TC treatment), both HIV-WT and HIV-45G viruses carried more GG-to-AG than GA-to-AA mutations but the differences were only significant for the HIV-45G infected animals (p=0.0020, Fig. 5A). At Day 58 post infection in treated mice (Fig. 5C), GG-to-AG rates were higher than GA-to-AA rates in both HIV-WT (p=0.0078) and HIV-45G (p=0.0020) infected mice. GG-to-AG mutation rates in HIV-WT and HIV-45G mice were comparable (p=0.6965) and were not due to selection of the RT-184I as rates were still comparable after analyses of sequences with codon 184 excluded (p=0.3445, Mann-Whitney).

**Fig. 5:**
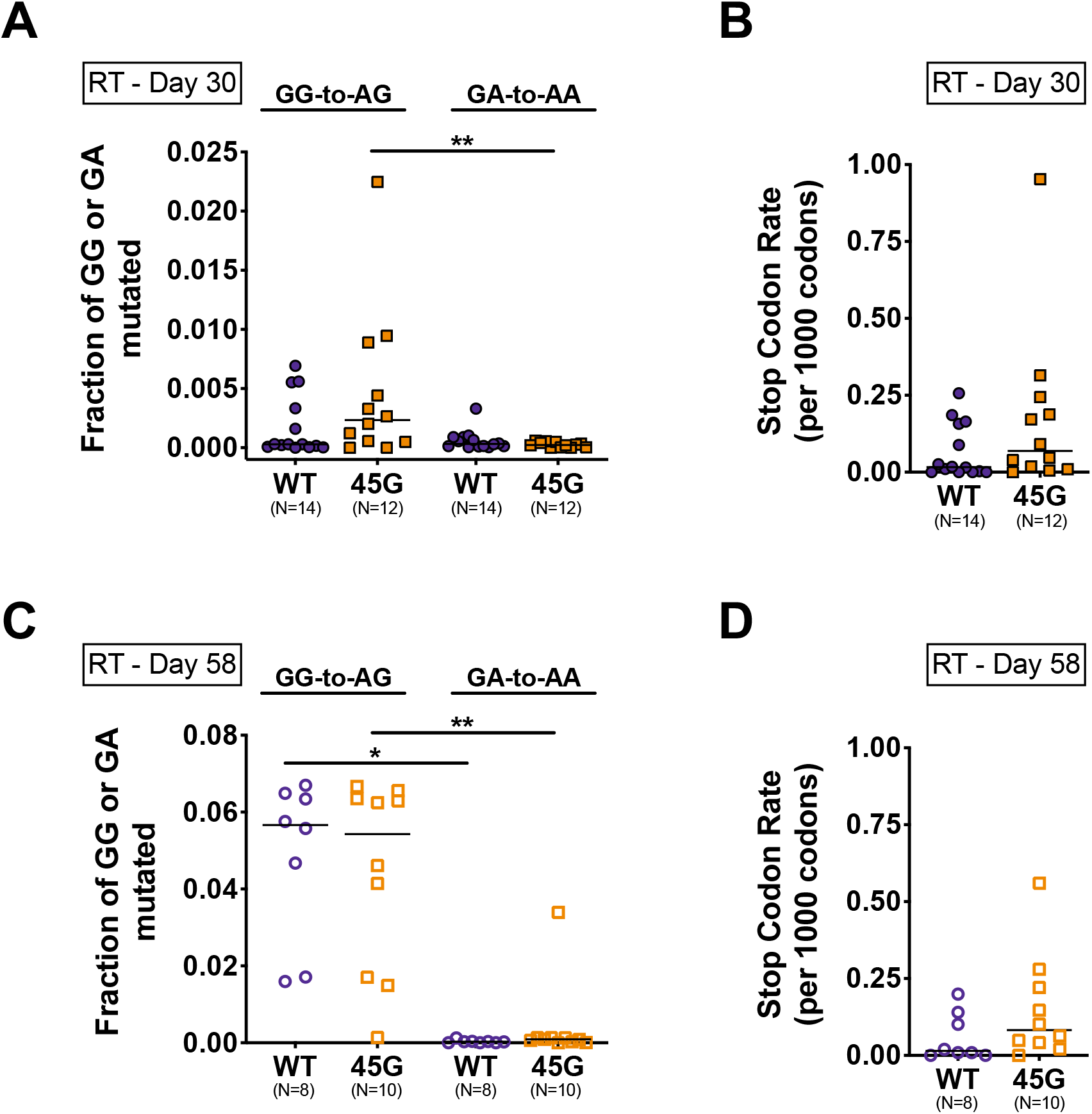
Mutagenesis in HIV *RT* gene in plasma viruses of HIV infected mice. **(A)** Dot plots depicting fractions of GG and GA dinucleotides mutated to AG (GG-to-AG) and AA (GA-to-AA), respectively, in individual mice (points) prior to 3TC treatment (Day 30 post infection). Bars depict means. p = 0.0020, Wilcoxon matched pairs signed rank test. **(B)** Stop codons were quantitated across all plasma sequences and a stop codon rate for every 1,000 codons sequenced was calculated for infected mice at Day 30 post infection. **(C)** Fraction of GG-to-AG or GA-to-AA dinucleotides mutations in plasma samples at Day 58 post infection in 3TC treated mice. (*) p ≤ 0.05, (**) p ≤ 0.01, Wilcoxon matched pairs signed rank test. **(D)** Stop codon rate in plasma samples from Day 58 post infection.

Due to its preferred dinucleotide context, A3G-induced mutagenesis can introduce mutations resulting in premature stop codons (e.g., UGG-to-UAG, (7, 69, 70)). Given that premature stop codons within RT are deleterious to replication (71–76), we were surprised to see that most infected animals (10/14 HIV-WT, 11/12 HIV-45G) had, at least, one plasma viral genome with mutations resulting in a stop codon upon protein translation. HIV-WT and HIV-45G viral populations in the plasma displayed stop codons predominantly at the four tryptophan (UGG-to-UAG or -UGA) codons (e.g., W212, W229, W239, W252) and rarely, at one of the six glutamine encoding codons (CAG-to-UAG, CAA-to-UAA; Q182, Q197, Q207, Q222, Q242, Q258). Of the 155,462 unique RT analyzed, 868 carried mutations leading to premature stop codon. The majority of the RT sequences (85%) only had a single stop codon but a small portion of sequences had two (11%), three (3%) or, at most, four (1%) stop codons. The rate of stop codons at Day 30 and at Day 58 post infection was, however, comparable between the two groups (p=0.1294, Fig. 5B; p=0.1211, Fig. 5D).

### Characterization of *vif* sequence diversity throughout the course of infection

Lastly, we were interested in the extent to which the *vif* sequences changed over the course of infection. We used samples remaining from a subset of 14 animals included in Experiment #3 to sequence a 268 bp long region of HIV *vif* (corresponding with codons 23 to 112) using a sequencing approach comprising UMID and Illumina 150 bp pair-end sequencing technology analogous to the approach taken for analyzing *RT* sequence diversity. In total, we generated 579,466 high quality paired-end reads representing 6,983 distinct UMIDs from 27 plasma vRNA samples.

We first looked for evidence of HIV-45G revertants at Day 30 and Day 58 post infection in five Vif-45G infected animals (Fig. 6A). Viruses in HIV-45G infected animals carrying the Vif-45E reversion were rare at Day 30 post infection (less than 1% in all five animals) but ranged between 0.5% and 9% of the plasma virus population present at Day 58 post infection (Fig. 6A). These data suggest that the Vif-45G genotype is quite stable over time.

**Fig. 6:**
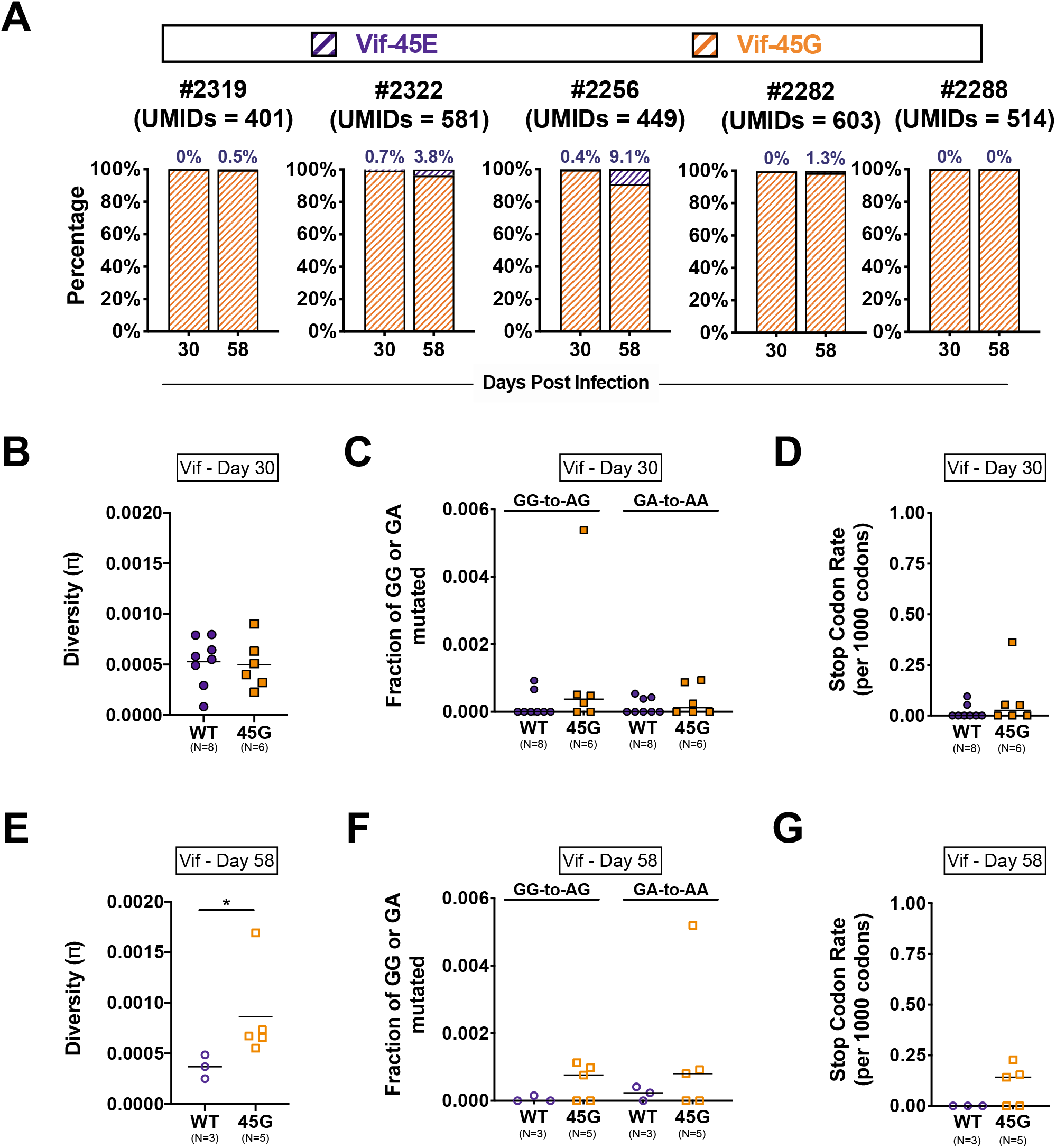
Characterizing HIV *Vif* mutations in infected mice. **(A)** Genotype of Vif codon 45 in five mice infected with HIV-45G and treated with 3TC from 30 to 58 Days post infection. Percentage of 45G and revertant E45 sequences indicated on stacked barplots. Percentages of the latter are annotated, as well. **(B)** Dot plots depicting nucleotide diversity (π) in *Vif* sequences in individual mice (points) infected with HIV-WT or HIV-45G at 30 Days post infection (prior to 3TC treatment). Bars depict means. **(C)** Dot plots depicting fractions of GG and GA dinucleotides mutated to AG (GG-to-AG) and AA (GA-to-AA), respectively, in individual mice (points) at 30 Days post infection. Bars depict means. **(D)** Stop codon rates were quantitated across plasma *Vif* sequences at Day 30 post infection (as in Fig. 5B). **(E)** Dot plots depicting π in *Vif* sequences in individual mice (points) at Day 58 post infection in 3TC treated mice. p = 0.0357, Mann-Whitney test. **(F)** Fraction of GG-to-AG or GA-to-AA dinucleotides mutated in *Vif* at Day 58 post infection in 3TC treated mice. **(G)** Stop codon rate in *Vif* sequences determined at Day 58 post infection in 3TC treated mice.

Since it has been suggested that *vif* diversification may be distinct from that of other HIV genes depending on the selective pressures exerted (7, 70, 77), we analyzed the nucleotide diversity, APOBEC3 driven mutagenesis and the rate of stop codons for the *vif* region sequenced (Figs. 6B-6G). The 268 bp long region of *vif* that we sequenced contains 17 or 18 GG (HIV-WT, Vif E45 = GAA; HIV-45G, Vif E45G = GGA) and 22 GA dinucleotides that could serve as APOBEC3 target motifs. Prior to treatment (Day 30 post infection), *vif* sequence diversity (Fig. 6B), GG-to-AG/GA-to-AA mutation rates (Fig. 6C) and stop codon frequency (Fig. 6C) were comparable between HIV-WT and HIV-45G infected mice. However, at Day 58 post infection, the Vif diversity was significantly higher in HIV-45G viruses compared to their wild-type counterparts (p=0.0357, Fig. 6E). APOBEC3 mutagenesis (GG-to-AG, GA-to-AA or rate of stop codon, Figs. 6F-6G) were comparable between the two groups.

Taken together, HIV-45G genotype can revert to wild-type Vif, but it only accounts for a small percentage of the circulating plasma variants in a portion of the mice tested. Moreover, there is some evidence suggesting that *RT* and *vif* regions diversity is caused by different mechanisms.

## DISCUSSION

Proviruses with footprints of past cytidine deamination are found in many, if not all, HIV infected patients (11, 17, 22, 60, 78–83). Nonetheless, it remains controversial to what extent APOBEC3-driven mutagenesis contributes to viral evolution and HIV/AIDS disease outcome *in vivo*. Some clinical studies find correlations between frequency of G-to-A mutations in proviruses and plasma viral loads (7, 82, 84), whereas others fail to find such associations (17, 83, 85). Controlled experiments in cell culture, however, provide strong experimental evidence in support of the notion that A3G-driven mutagenesis facilitates HIV diversification and promote escape from selection pressure (44, 69, 86). In the current study, we perform controlled, *in vivo* infection experiments to provide new insights into the dynamics of HIV diversification within the plasma compartment of humanized mice in the absence and presence of selection pressure. We show that suboptimal neutralization of A3G shapes the phenotype of circulating viruses and compromises 3TC treatment outcomes.

Previous studies in cell culture (20, 44, 87) and in humanized mice (33–37) have examined the role of APOBEC3 proteins in HIV replication and pathogenesis *in vivo* but our study dissects the impact of A3G-driven mutagenesis in the context of viral evolution by introducing selection in the form of antiretroviral treatment. Moreover, we focus on viral diversification within the plasma compartment, which reflects the actively replicating viral quasispecies in an immediate and dynamic manner. HIV-45G, which has suboptimal anti-A3G activity, is less fit than HIV-WT in treatment-naïve animals overtime (Fig. 2A) pointing to the slow accumulation of mutations that fail to provide any evolutionary benefit given that hu-NSG mice lack immunologic pressures (i.e., CD4+/CD8+ T cell responses or antibodies (88)). Conversely, HIV-45G responds less well and rebounds more rapidly in the presence of 3TC treatment (Figs. 2B-2D). Thus, complete inactivation of A3G is dispensable for initiating a productive and robust infection of humanized mice and suboptimal A3G neutralization can be beneficial to HIV for overcoming evolutionary bottlenecks such as selection pressure by antiretroviral drugs.

Sequencing technologies have dramatically improved over the last decade allowing for high-resolution, accurate representation of viral quasispecies. We combined UMIDs with Illumina pair-end sequencing to analyze >150,000 individual *RT* sequences sampling approximately 1,170 distinct viral genomes for each individual time point. Previous studies in humanized mice examined sequence diversity using bulk amplification followed by sequencing of individual clones (33–36) or by single-genome sequencing (SGS, (36)). The first of these methods fails to reliably distinguish between individual variants and may skew viral diversity measurements due to PCR errors or bias (61, 62). SGS is regarded as gold standard in the field since it analyses distinct genomes and provides information on large regions. However, the approach is very work intensive and costly, limiting the numbers of genomes that can be sampled (89–91). For example, one previous study used SGS to analyze a total of 265 genomes from eight mice (36). In our study, in contrast, we analyzed on average 1,400 *RT* sequences per infected animal providing us with solid data on minority viral populations. It has to be noted that even at our high sequencing depth, we only sample a limited portion of the viruses circulating in the plasma compartment (i.e., average plasma viremia at Day 30 post infection is 89,620 copies/mL). Despite this limitation, our sequencing data revealed a number of previously overlooked facts regarding the dynamics of HIV evolution *in vivo*. First, we found that 3TC drug resistant viral variants were found, in a third of the animals, prior to 3TC treatment initiation (Fig. 3B). Pre-existing 3TC resistance was, however, not linked to more rapid treatment failure suggesting that these viruses may not be replication competent. Second, we noted a positive correlation between increased *RT* sequence diversity and high plasma viremia in HIV-45G infected animals after 30 days of unchecked replication (Fig. 4B). This association is surprising since conventional wisdom would predict the opposite to be true. Indeed, this is exactly what we observe for HIV-WT infections where increased *RT* diversity is associated with lower plasma viremia (Fig. 4C). Third, the kinetics with which the two drug resistant variants RT-184I and RT-184V appeared in the plasma differed between the two viruses. In mice infected with HIV-45G, the RT-184I variants arose at a rate 8.3-times faster than that of RT-184V variants, whereas emergence rates for these variants were comparable in mice infected with HIV-WT (Fig. 3D). However, RT-184I did not appear more readily in HIV-45G infected animals. This could be due to the fact that our earliest collection time point was ten days after treatment initiation. It is also conceivable that drug resistant variants first evolve, replicate and expand in tissue compartments with the plasma compartment being a mere reflection after the fact.

Taken together, future studies investigating the viral diversification at the viral RNA, cellular viral RNA and proviral level in vivo in the humanized model will provide further insights into how viral quasispecies shaped by APOBEC3 mutagenesis inform on HIV pathogenesis.

## ACKNOWLEDGMENTS

We thank the Speck, Sachidanandam and Simon laboratories for insightful discussions. This work was funded in part by NIH/NIAID grants AI064001, AI120998 (VS); NIH/NIGMS grant GM113886 (LCF), NIH/NIGMS grant T32-GM007280 (MMH), the pre- and post-doctoral USPHS Institutional Research Training Award T32-AI07647 (MMH), the clinical research focus program “Human Hemato-Lymphatic Diseases” of the University of Zurich (RFS) and SNF #310031_153248/1 and matching funds, University of Zurich (RFS).

---

**Supplemental Table 1.**
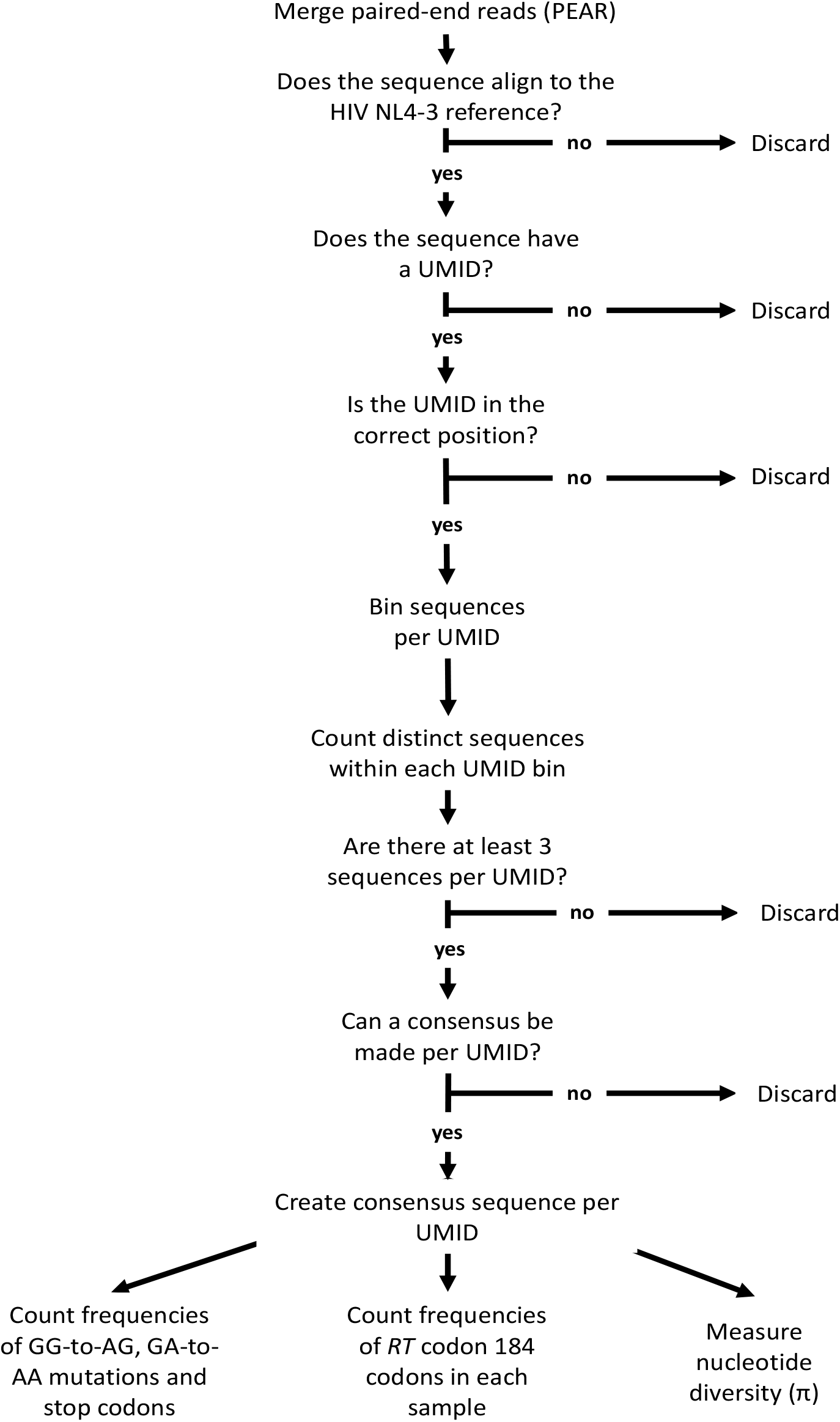
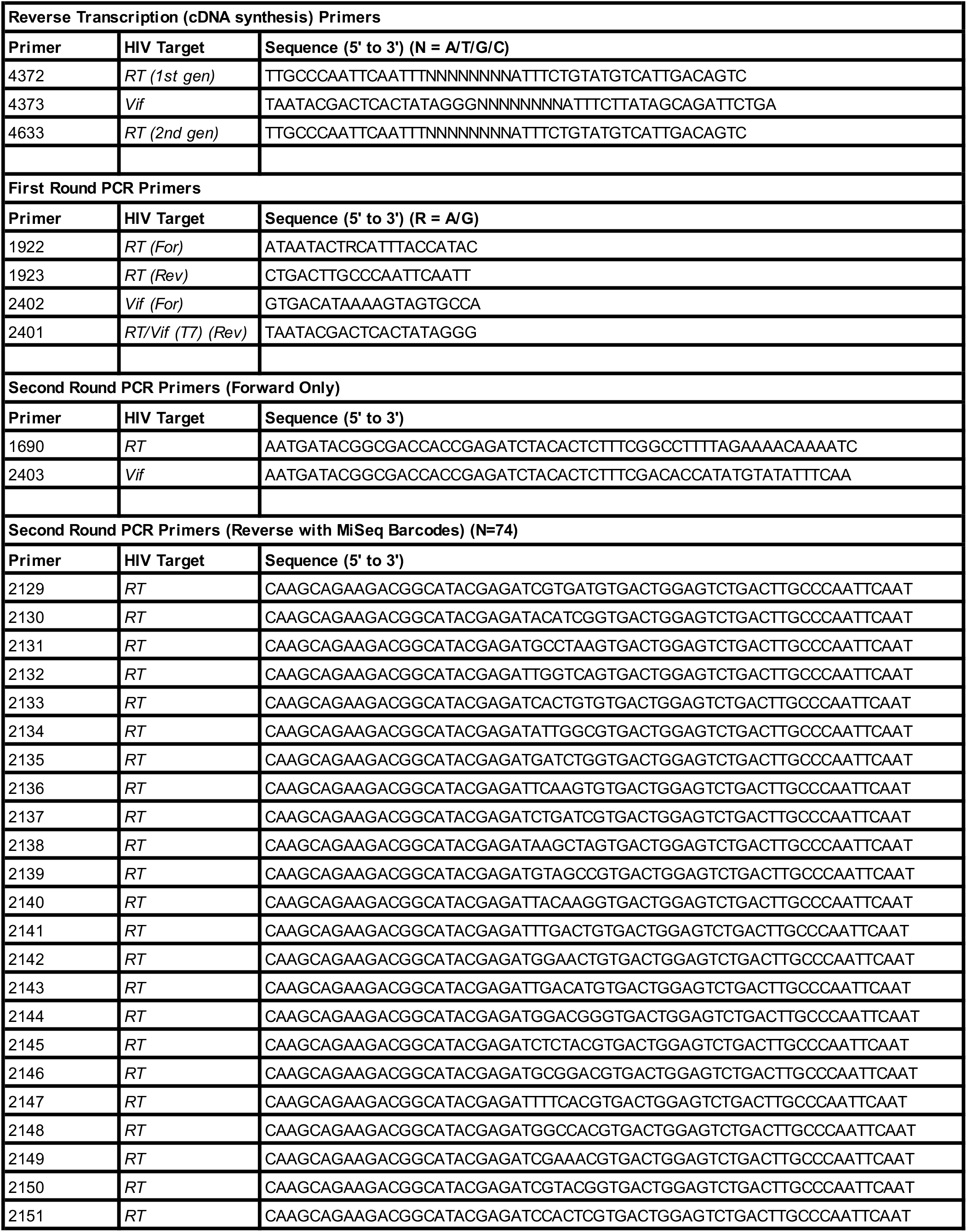
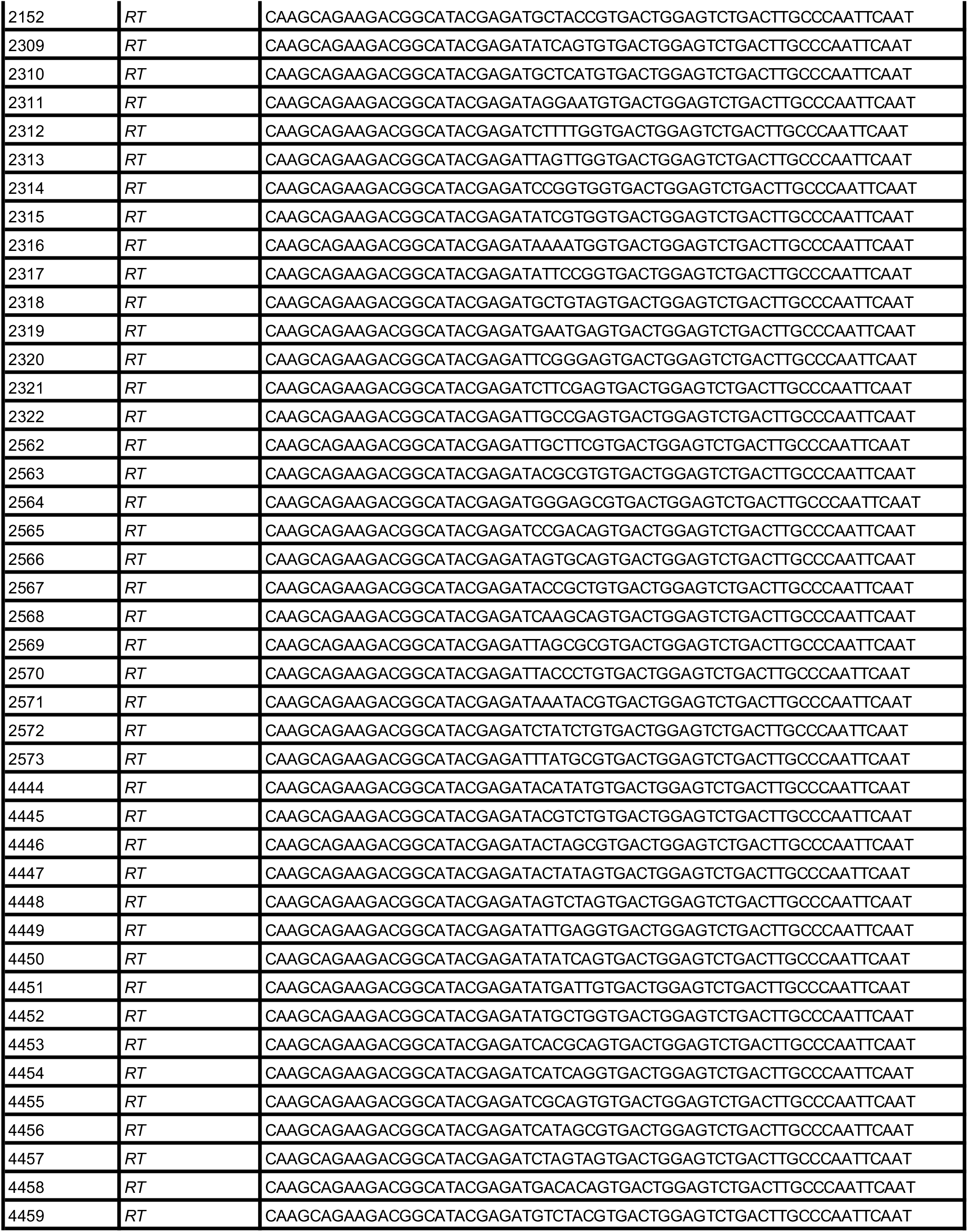
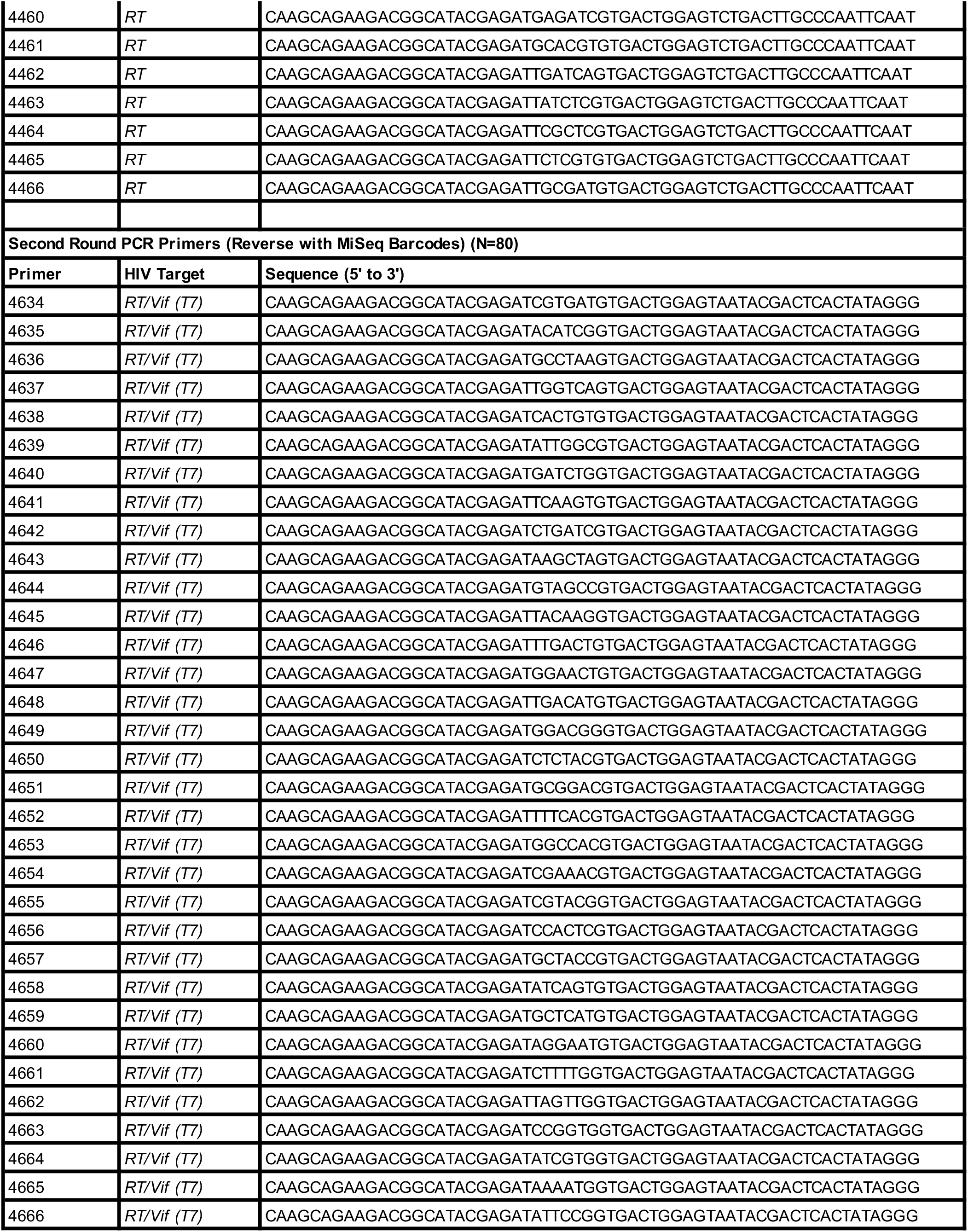
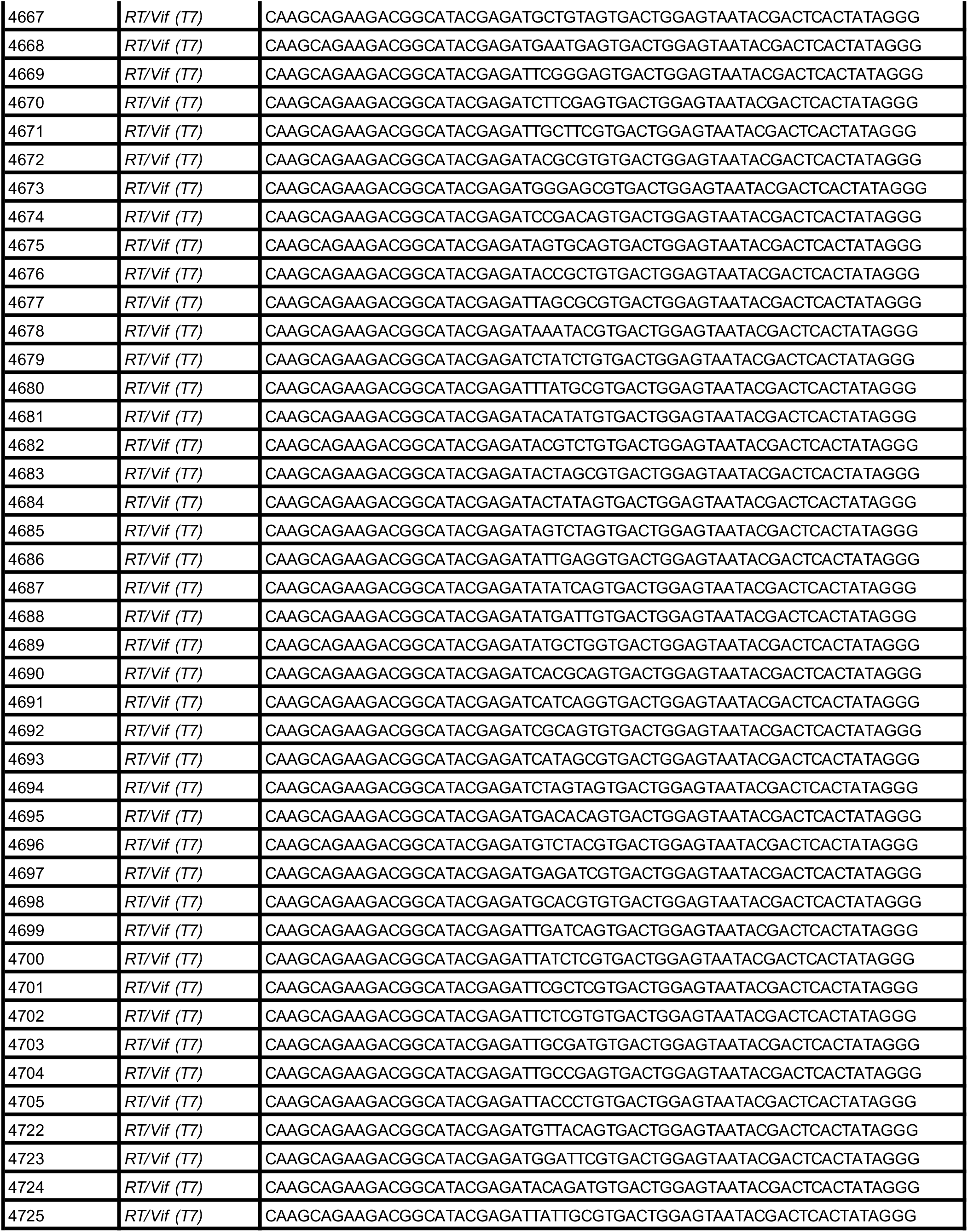
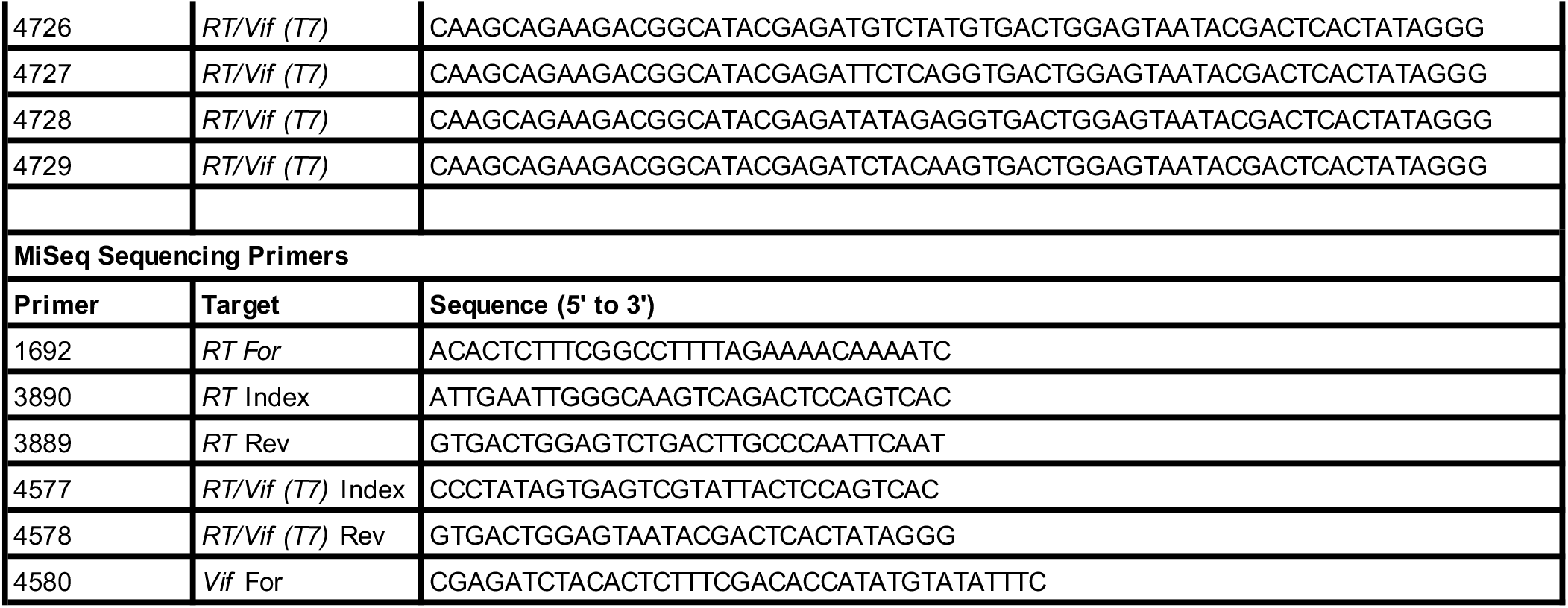
Sequencing pipeline and primers for reverse transcription, PCR amplification and Illumina MiSeq sequencing.

## REFERENCES

1. Wood N, Bhattacharya T, Keele BF, Giorgi E, Liu M, Gaschen B, Daniels M, Ferrari G, Haynes BF, McMichael A, Shaw GM, Hahn BH, Korber B, Seoighe C. 2009. HIV evolution in early infection: selection pressures, patterns of insertion and deletion, and the impact of APOBEC. PLoS Pathog 5:e1000414.

2. Rhodes TD, Nikolaitchik O, Chen J, Powell D, Hu WS. 2005. Genetic recombination of human immunodeficiency virus type 1 in one round of viral replication: effects of genetic distance, target cells, accessory genes, and lack of high negative interference in crossover events. J Virol 79:1666–77.

3. Smyth RP, Negroni M. 2016. A step forward understanding HIV-1 diversity. Retrovirology 13:27.

4. Ji JP, Loeb LA. 1992. Fidelity of HIV-1 reverse transcriptase copying RNA in vitro. Biochemistry 31:954–8.

5. Hu WS, Hughes SH. 2012. HIV-1 reverse transcription. Cold Spring Harb Perspect Med 2.

6. Malim MH. 2009. APOBEC proteins and intrinsic resistance to HIV-1 infection. Philos Trans R Soc Lond B Biol Sci 364:675–87.

7. Cuevas JM, Geller R, Garijo R, Lopez-Aldeguer J, Sanjuan R. 2015. Extremely High Mutation Rate of HIV-1 In Vivo. PLoS Biol 13:e1002251.

8. van Zyl G, Bale MJ, Kearney MF. 2018. HIV evolution and diversity in ART-treated patients. Retrovirology 15:14.

9. Desimmie BA, Delviks-Frankenberrry KA, Burdick RC, Qi D, Izumi T, Pathak VK. 2014. Multiple APOBEC3 restriction factors for HIV-1 and one Vif to rule them all. J Mol Biol 426:1220–45.

10. Simon V, Bloch N, Landau NR. 2015. Intrinsic host restrictions to HIV-1 and mechanisms of viral escape. Nat Immunol 16:546–53.

11. Armitage AE, Deforche K, Welch JJ, Van Laethem K, Camacho R, Rambaut A, Iversen AK. 2014. Possible footprints of APOBEC3F and/or other APOBEC3 deaminases, but not APOBEC3G, on HIV-1 from patients with acute/early and chronic infections. J Virol 88:12882–94.

12. Chaipan C, Smith JL, Hu WS, Pathak VK. 2013. APOBEC3G restricts HIV-1 to a greater extent than APOBEC3F and APOBEC3DE in human primary CD4+ T cells and macrophages. J Virol 87:444–53.

13. Harris RS, Bishop KN, Sheehy AM, Craig HM, Petersen-Mahrt SK, Watt IN, Neuberger MS, Malim MH. 2003. DNA deamination mediates innate immunity to retroviral infection. Cell 113:803–9.

14. Albin JS, Harris RS. 2010. Interactions of host APOBEC3 restriction factors with HIV-1 in vivo: implications for therapeutics. Expert Rev Mol Med 12:e4.

15. Janini M, Rogers M, Birx DR, McCutchan FE. 2001. Human Immunodeficiency Virus Type 1 DNA Sequences Genetically Damaged by Hypermutation Are Often Abundant in Patient Peripheral Blood Mononuclear Cells and May Be Generated during Near-Simultaneous Infection and Activation of CD4+ T Cells. Journal of Virology 75:7973–7986.

16. Russell RA, Moore MD, Hu WS, Pathak VK. 2009. APOBEC3G induces a hypermutation gradient: purifying selection at multiple steps during HIV-1 replication results in levels of G-to-A mutations that are high in DNA, intermediate in cellular viral RNA, and low in virion RNA. Retrovirology 6:16.

17. Gandhi SK, Siliciano JD, Bailey JR, Siliciano RF, Blankson JN. 2008. Role of APOBEC3G/F-mediated hypermutation in the control of human immunodeficiency virus type 1 in elite suppressors. J Virol 82:3125–30.

18. Simon V, Zennou V, Murray D, Huang Y, Ho DD, Bieniasz PD. 2005. Natural variation in Vif: differential impact on APOBEC3G/3F and a potential role in HIV-1 diversification. PLoS Pathog 1:e6.

19. Reddy K, Ooms M, Letko M, Garrett N, Simon V, Ndung’u T. 2016. Functional characterization of Vif proteins from HIV-1 infected patients with different APOBEC3G haplotypes. AIDS 30:1723–9.

20. Fourati S, Malet I, Binka M, Boukobza S, Wirden M, Sayon S, Simon A, Katlama C, Simon V, Calvez V, Marcelin AG. 2010. Partially active HIV-1 Vif alleles facilitate viral escape from specific antiretrovirals. AIDS 24:2313–21.

21. Kourteva Y, De Pasquale M, Allos T, McMunn C, D’Aquila RT. 2012. APOBEC3G expression and hypermutation are inversely associated with human immunodeficiency virus type 1 (HIV-1) burden in vivo. Virology 430:1–9.

22. Kim EY, Lorenzo-Redondo R, Little SJ, Chung YS, Phalora PK, Maljkovic Berry I, Archer J, Penugonda S, Fischer W, Richman DD, Bhattacharya T, Malim MH, Wolinsky SM. 2014. Human APOBEC3 induced mutation of human immunodeficiency virus type-1 contributes to adaptation and evolution in natural infection. PLoS Pathog 10:e1004281.

23. Shultz LD, Brehm MA, Garcia-Martinez JV, Greiner DL. 2012. Humanized mice for immune system investigation: progress, promise and challenges. Nat Rev Immunol 12:786–98.

24. Dudek TE, No DC, Seung E, Vrbanac VD, Fadda L, Bhoumik P, Boutwell CL, Power KA, Gladden AD, Battis L, Mellors EF, Tivey TR, Gao X, Altfeld M, Luster AD, Tager AM, Allen TM. 2012. Rapid evolution of HIV-1 to functional CD8(+) T cell responses in humanized BLT mice. Sci Transl Med 4:143ra98.

25. Melkus MW, Estes JD, Padgett-Thomas A, Gatlin J, Denton PW, Othieno FA, Wege AK, Haase AT, Garcia JV. 2006. Humanized mice mount specific adaptive and innate immune responses to EBV and TSST-1. Nat Med 12:1316–22.

26. Denton PW, Olesen R, Choudhary SK, Archin NM, Wahl A, Swanson MD, Chateau M, Nochi T, Krisko JF, Spagnuolo RA, Margolis DM, Garcia JV. 2012. Generation of HIV latency in humanized BLT mice. J Virol 86:630–4.

27. Denton PW, Othieno F, Martinez-Torres F, Zou W, Krisko JF, Fleming E, Zein S, Powell DA, Wahl A, Kwak YT, Welch BD, Kay MS, Payne DA, Gallay P, Appella E, Estes JD, Lu M, Garcia JV. 2011. One percent tenofovir applied topically to humanized BLT mice and used according to the CAPRISA 004 experimental design demonstrates partial protection from vaginal HIV infection, validating the BLT model for evaluation of new microbicide candidates. J Virol 85:7582–93.

28. Horwitz JA, Halper-Stromberg A, Mouquet H, Gitlin AD, Tretiakova A, Eisenreich TR, Malbec M, Gravemann S, Billerbeck E, Dorner M, Buning H, Schwartz O, Knops E, Kaiser R, Seaman MS, Wilson JM, Rice CM, Ploss A, Bjorkman PJ, Klein F, Nussenzweig MC. 2013. HIV-1 suppression and durable control by combining single broadly neutralizing antibodies and antiretroviral drugs in humanized mice. Proc Natl Acad Sci U S A 110:16538–43.

29. Nischang M, Sutmuller R, Gers-Huber G, Audige A, Li D, Rochat MA, Baenziger S, Hofer U, Schlaepfer E, Regenass S, Amssoms K, Stoops B, Van Cauwenberge A, Boden D, Kraus G, Speck RF. 2012. Humanized mice recapitulate key features of HIV-1 infection: a novel concept using long-acting anti-retroviral drugs for treating HIV-1. PLoS One 7:e38853.

30. Ince WL, Zhang L, Jiang Q, Arrildt K, Su L, Swanstrom R. 2010. Evolution of the HIV-1 env gene in the Rag2-/- gammaC-/- humanized mouse model. J Virol 84:2740–52.

31. Yamada E, Yoshikawa R, Nakano Y, Misawa N, Koyanagi Y, Sato K. 2015. Impacts of humanized mouse models on the investigation of HIV-1 infection: illuminating the roles of viral accessory proteins in vivo. Viruses 7:1373–90.

32. Zhang L, Su L. 2012. HIV-1 immunopathogenesis in humanized mouse models. Cell Mol Immunol 9:237–44.

33. Sato K, Izumi T, Misawa N, Kobayashi T, Yamashita Y, Ohmichi M, Ito M, Takaori-Kondo A, Koyanagi Y. 2010. Remarkable lethal G-to-A mutations in vif-proficient HIV-1 provirus by individual APOBEC3 proteins in humanized mice. J Virol 84:9546–56.

34. Krisko JF, Martinez-Torres F, Foster JL, Garcia JV. 2013. HIV restriction by APOBEC3 in humanized mice. PLoS Pathog 9:e1003242.

35. Krisko JF, Begum N, Baker CE, Foster JL, Garcia JV. 2016. APOBEC3G and APOBEC3F Act in Concert To Extinguish HIV-1 Replication. J Virol 90:4681–4695.

36. Sato K, Takeuchi JS, Misawa N, Izumi T, Kobayashi T, Kimura Y, Iwami S, Takaori-Kondo A, Hu WS, Aihara K, Ito M, An DS, Pathak VK, Koyanagi Y. 2014. APOBEC3D and APOBEC3F potently promote HIV-1 diversification and evolution in humanized mouse model. PLoS Pathog 10:e1004453.

37. Nakano Y, Misawa N, Juarez-Fernandez G, Moriwaki M, Nakaoka S, Funo T, Yamada E, Soper A, Yoshikawa R, Ebrahimi D, Tachiki Y, Iwami S, Harris RS, Koyanagi Y, Sato K. 2017. HIV-1 competition experiments in humanized mice show that APOBEC3H imposes selective pressure and promotes virus adaptation. PLoS Pathog 13:e1006348.

38. Derdeyn CA, Decker JM, Sfakianos JN, Wu X, O’Brien WA, Ratner L, Kappes JC, Shaw GM, Hunter E. 2000. Sensitivity of human immunodeficiency virus type 1 to the fusion inhibitor T-20 is modulated by coreceptor specificity defined by the V3 loop of gp120. J Virol 74:8358–67.

39. Platt EJ, Bilska M, Kozak SL, Kabat D, Montefiori DC. 2009. Evidence that ecotropic murine leukemia virus contamination in TZM-bl cells does not affect the outcome of neutralizing antibody assays with human immunodeficiency virus type 1. J Virol 83:8289–92.

40. Platt EJ, Wehrly K, Kuhmann SE, Chesebro B, Kabat D. 1998. Effects of CCR5 and CD4 cell surface concentrations on infections by macrophagetropic isolates of human immunodeficiency virus type 1. J Virol 72:2855–64.

41. Takeuchi Y, McClure MO, Pizzato M. 2008. Identification of gammaretroviruses constitutively released from cell lines used for human immunodeficiency virus research. J Virol 82:12585–8.

42. Wei X, Decker JM, Liu H, Zhang Z, Arani RB, Kilby JM, Saag MS, Wu X, Shaw GM, Kappes JC. 2002. Emergence of resistant human immunodeficiency virus type 1 in patients receiving fusion inhibitor (T-20) monotherapy. Antimicrob Agents Chemother 46:1896–905.

43. Adachi A, Gendelman HE, Koenig S, Folks T, Willey R, Rabson A, Martin MA. 1986. Production of acquired immunodeficiency syndrome-associated retrovirus in human and nonhuman cells transfected with an infectious molecular clone. J Virol 59:284–91.

44. Mulder LC, Harari A, Simon V. 2008. Cytidine deamination induced HIV-1 drug resistance. Proc Natl Acad Sci U S A 105:5501–6.

45. Moore JP, McKeating JA, Weiss RA, Sattentau QJ. 1990. Dissociation of gp120 from HIV-1 virions induced by soluble CD4. Science 250:1139–42.

46. Zhang J, Kobert K, Flouri T, Stamatakis A. 2014. PEAR: a fast and accurate Illumina Paired-End reAd mergeR. Bioinformatics 30:614–20.

47. Librado P, Rozas J. 2009. DnaSP v5: a software for comprehensive analysis of DNA polymorphism data. Bioinformatics 25:1451–2.

48. Ooms M, Brayton B, Letko M, Maio SM, Pilcher CD, Hecht FM, Barbour JD, Simon V. 2013. HIV-1 Vif adaptation to human APOBEC3H haplotypes. Cell Host Microbe 14:411–21.

49. Russell RA, Pathak VK. 2007. Identification of two distinct human immunodeficiency virus type 1 Vif determinants critical for interactions with human APOBEC3G and APOBEC3F. J Virol 81:8201–10.

50. Feng Y, Baig TT, Love RP, Chelico L. 2014. Suppression of APOBEC3-mediated restriction of HIV-1 by Vif. Front Microbiol 5:450.

51. Henriet S, Mercenne G, Bernacchi S, Paillart JC, Marquet R. 2009. Tumultuous relationship between the human immunodeficiency virus type 1 viral infectivity factor (Vif) and the human APOBEC-3G and APOBEC-3F restriction factors. Microbiol Mol Biol Rev 73:211–32.

52. Schinazi RF, Lloyd RM, Jr., Nguyen MH, Cannon DL, McMillan A, Ilksoy N, Chu CK, Liotta DC, Bazmi HZ, Mellors JW. 1993. Characterization of human immunodeficiency viruses resistant to oxathiolane-cytosine nucleosides. Antimicrob Agents Chemother 37:875–81.

53. Schuurman R, Nijhuis M, van Leeuwen R, Schipper P, de Jong D, Collis P, Danner SA, Mulder J, Loveday C, Christopherson C, et al. 1995. Rapid changes in human immunodeficiency virus type 1 RNA load and appearance of drug-resistant virus populations in persons treated with lamivudine (3TC). J Infect Dis 171:1411–9.

54. Wainberg MA, Drosopoulos WC, Salomon H, Hsu M, Borkow G, Parniak M, Gu Z, Song Q, Manne J, Islam S, Castriota G, Prasad VR. 1996. Enhanced fidelity of 3TC-selected mutant HIV-1 reverse transcriptase. Science 271:1282–5.

55. Keulen W, Back NK, van Wijk A, Boucher CA, Berkhout B. 1997. Initial appearance of the 184Ile variant in lamivudine-treated patients is caused by the mutational bias of human immunodeficiency virus type 1 reverse transcriptase. J Virol 71:3346–50.

56. Sarafianos SG, Das K, Clark AD, Jr., Ding J, Boyer PL, Hughes SH, Arnold E. 1999. Lamivudine (3TC) resistance in HIV-1 reverse transcriptase involves steric hindrance with beta-branched amino acids. Proc Natl Acad Sci U S A 96:10027–32.

57. Gao HQ, Boyer PL, Sarafianos SG, Arnold E, Hughes SH. 2000. The role of steric hindrance in 3TC resistance of human immunodeficiency virus type-1 reverse transcriptase. J Mol Biol 300:403–18.

58. Frost SD, Nijhuis M, Schuurman R, Boucher CA, Brown AJ. 2000. Evolution of lamivudine resistance in human immunodeficiency virus type 1-infected individuals: the relative roles of drift and selection. J Virol 74:6262–8.

59. Brenner BG, Turner D, Wainberg MA. 2002. HIV-1 drug resistance: can we overcome? Expert Opin Biol Ther 2:751–61.

60. Berkhout B, de Ronde A. 2004. APOBEC3G versus reverse transcriptase in the generation of HIV-1 drug-resistance mutations. AIDS 18:1861–3.

61. Jabara CB, Jones CD, Roach J, Anderson JA, Swanstrom R. 2011. Accurate sampling and deep sequencing of the HIV-1 protease gene using a Primer ID. Proc Natl Acad Sci U S A 108:20166–71.

62. Keys JR, Zhou S, Anderson JA, Eron JJ, Jr., Rackoff LA, Jabara C, Swanstrom R. 2015. Primer ID Informs Next-Generation Sequencing Platforms and Reveals Preexisting Drug Resistance Mutations in the HIV-1 Reverse Transcriptase Coding Domain. AIDS Res Hum Retroviruses 31:658–68.

63. Nei M, Kumar, S. 2000. Molecular Evolution and Phylogenetics. Oxford University Press, New York.

64. Nei M, Gojobori T. 1986. Simple methods for estimating the numbers of synonymous and nonsynonymous nucleotide substitutions. Mol Biol Evol 3:418–26.

65. Nelson CW, Hughes AL. 2015. Within-host nucleotide diversity of virus populations: insights from next-generation sequencing. Infect Genet Evol 30:1–7.

66. Alizon S, Magnus C. 2012. Modelling the course of an HIV infection: insights from ecology and evolution. Viruses 4:1984–2013.

67. Lima K, Leal E, Cavalcanti AMS, Salustiano DM, de Medeiros LB, da Silva SP, Lacerda HR. 2017. Increase in human immunodeficiency virus 1 diversity and detection of various subtypes and recombinants in north-eastern Brazil. J Med Microbiol 66:526–535.

68. Refsland EW, Hultquist JF, Harris RS. 2012. Endogenous origins of HIV-1 G-to-A hypermutation and restriction in the nonpermissive T cell line CEM2n. PLoS Pathog 8:e1002800.

69. Sadler HA, Stenglein MD, Harris RS, Mansky LM. 2010. APOBEC3G contributes to HIV-1 variation through sublethal mutagenesis. J Virol 84:7396–404.

70. Armitage AE, Deforche K, Chang CH, Wee E, Kramer B, Welch JJ, Gerstoft J, Fugger L, McMichael A, Rambaut A, Iversen AK. 2012. APOBEC3G-induced hypermutation of human immunodeficiency virus type-1 is typically a discrete “all or nothing” phenomenon. PLoS Genet 8:e1002550.

71. Bruner KM, Murray AJ, Pollack RA, Soliman MG, Laskey SB, Capoferri AA, Lai J, Strain MC, Lada SM, Hoh R, Ho YC, Richman DD, Deeks SG, Siliciano JD, Siliciano RF. 2016. Defective proviruses rapidly accumulate during acute HIV-1 infection. Nat Med 22:1043–9.

72. Ho YC, Shan L, Hosmane NN, Wang J, Laskey SB, Rosenbloom DI, Lai J, Blankson JN, Siliciano JD, Siliciano RF. 2013. Replication-competent noninduced proviruses in the latent reservoir increase barrier to HIV-1 cure. Cell 155:540–51.

73. Imamichi H, Dewar RL, Adelsberger JW, Rehm CA, O’Doherty U, Paxinos EE, Fauci AS, Lane HC. 2016. Defective HIV-1 proviruses produce novel protein-coding RNA species in HIV-infected patients on combination antiretroviral therapy. Proc Natl Acad Sci U S A 113:8783–8.

74. Pollack RA, Jones RB, Pertea M, Bruner KM, Martin AR, Thomas AS, Capoferri AA, Beg SA, Huang SH, Karandish S, Hao H, Halper-Stromberg E, Yong PC, Kovacs C, Benko E, Siliciano RF, Ho YC. 2017. Defective HIV-1 Proviruses Are Expressed and Can Be Recognized by Cytotoxic T Lymphocytes, which Shape the Proviral Landscape. Cell Host Microbe 21:494–506 e4.

75. Grant RM, Abrams DI. 1998. Not all is dead in HIV-1 graveyard. Lancet 351:308–9.

76. Maldarelli F. 2016. The role of HIV integration in viral persistence: no more whistling past the proviral graveyard. J Clin Invest 126:438–47.

77. Kijak GH, Janini LM, Tovanabutra S, Sanders-Buell E, Arroyo MA, Robb ML, Michael NL, Birx DL, McCutchan FE. 2008. Variable contexts and levels of hypermutation in HIV-1 proviral genomes recovered from primary peripheral blood mononuclear cells. Virology 376:101–11.

78. Fourati S, Lambert-Niclot S, Soulie C, Malet I, Valantin MA, Descours B, Ait-Arkoub Z, Mory B, Carcelain G, Katlama C, Calvez V, Marcelin AG. 2012. HIV-1 genome is often defective in PBMCs and rectal tissues after long-term HAART as a result of APOBEC3 editing and correlates with the size of reservoirs. J Antimicrob Chemother 67:2323–6.

79. Jern P, Russell RA, Pathak VK, Coffin JM. 2009. Likely role of APOBEC3G-mediated G-to-A mutations in HIV-1 evolution and drug resistance. PLoS Pathog 5:e1000367.

80. Karlsson AC, Iversen AK, Chapman JM, de Oliviera T, Spotts G, McMichael AJ, Davenport MP, Hecht FM, Nixon DF. 2007. Sequential broadening of CTL responses in early HIV-1 infection is associated with viral escape. PLoS One 2:e225.

81. Kieffer TL, Finucane MM, Nettles RE, Quinn TC, Broman KW, Ray SC, Persaud D, Siliciano RF. 2004. Genotypic analysis of HIV-1 drug resistance at the limit of detection: virus production without evolution in treated adults with undetectable HIV loads. J Infect Dis 189:1452–65.

82. Pace C, Keller J, Nolan D, James I, Gaudieri S, Moore C, Mallal S. 2006. Population level analysis of human immunodeficiency virus type 1 hypermutation and its relationship with APOBEC3G and vif genetic variation. J Virol 80:9259–69.

83. Piantadosi A, Humes D, Chohan B, McClelland RS, Overbaugh J. 2009. Analysis of the percentage of human immunodeficiency virus type 1 sequences that are hypermutated and markers of disease progression in a longitudinal cohort, including one individual with a partially defective Vif. J Virol 83:7805–14.

84. de Lima-Stein ML, Alkmim WT, Bizinoto MC, Lopez LF, Burattini MN, Maricato JT, Giron L, Sucupira MC, Diaz RS, Janini LM. 2014. In vivo HIV-1 hypermutation and viral loads among antiretroviral-naive Brazilian patients. AIDS Res Hum Retroviruses 30:867–80.

85. Amoedo ND, Afonso AO, Cunha SM, Oliveira RH, Machado ES, Soares MA. 2011. Expression of APOBEC3G/3F and G-to-A hypermutation levels in HIV-1-infected children with different profiles of disease progression. PLoS One 6:e24118.

86. Kim EY, Bhattacharya T, Kunstman K, Swantek P, Koning FA, Malim MH, Wolinsky SM. 2010. Human APOBEC3G-mediated editing can promote HIV-1 sequence diversification and accelerate adaptation to selective pressure. J Virol 84:10402–5.

87. Hache G, Abbink TE, Berkhout B, Harris RS. 2009. Optimal translation initiation enables Vif-deficient human immunodeficiency virus type 1 to escape restriction by APOBEC3G. J Virol 83:5956–60.

88. Nixon CC, Mavigner M, Silvestri G, Garcia JV. 2017. In Vivo Models of Human Immunodeficiency Virus Persistence and Cure Strategies. J Infect Dis 215:S142–S151.

89. Salazar-Gonzalez JF, Bailes E, Pham KT, Salazar MG, Guffey MB, Keele BF, Derdeyn CA, Farmer P, Hunter E, Allen S, Manigart O, Mulenga J, Anderson JA, Swanstrom R, Haynes BF, Athreya GS, Korber BT, Sharp PM, Shaw GM, Hahn BH. 2008. Deciphering human immunodeficiency virus type 1 transmission and early envelope diversification by single-genome amplification and sequencing. J Virol 82:3952–70.

90. Keele BF, Giorgi EE, Salazar-Gonzalez JF, Decker JM, Pham KT, Salazar MG, Sun C, Grayson T, Wang S, Li H, Wei X, Jiang C, Kirchherr JL, Gao F, Anderson JA, Ping LH, Swanstrom R, Tomaras GD, Blattner WA, Goepfert PA, Kilby JM, Saag MS, Delwart EL, Busch MP, Cohen MS, Montefiori DC, Haynes BF, Gaschen B, Athreya GS, Lee HY, Wood N, Seoighe C, Perelson AS, Bhattacharya T, Korber BT, Hahn BH, Shaw GM. 2008. Identification and characterization of transmitted and early founder virus envelopes in primary HIV-1 infection. Proc Natl Acad Sci U S A 105:7552–7.

91. Palmer S, Kearney M, Maldarelli F, Halvas EK, Bixby CJ, Bazmi H, Rock D, Falloon J, Davey RT, Jr., Dewar RL, Metcalf JA, Hammer S, Mellors JW, Coffin JM. 2005. Multiple, linked human immunodeficiency virus type 1 drug resistance mutations in treatment-experienced patients are missed by standard genotype analysis. J Clin Microbiol 43:406–13.

